# Do deep neural networks see the way we do?

**DOI:** 10.1101/860759

**Authors:** Georgin Jacob, R. T. Pramod, Harish Katti, S. P. Arun

**Affiliations:** Centre for Neuroscience, Indian Institute of Science, Bangalore 560012; Department of Electrical Communication Engineering, Indian Institute of Science, Bangalore 560012

## Abstract

Deep neural networks have revolutionized computer vision, and their object representations match coarsely with the brain. As a result, it is widely believed that any fine scale differences between deep networks and brains can be fixed with increased training data or minor changes in architecture. But what if there are qualitative differences between brains and deep networks? Do deep networks even see the way we do? To answer this question, we chose a deep neural network optimized for object recognition and asked whether it exhibits well-known perceptual and neural phenomena despite not being explicitly trained to do so. To our surprise, many phenomena were present in the network, including the Thatcher effect, mirror confusion, Weber’s law, relative size, multiple object normalization and sparse coding along multiple dimensions. However, some perceptual phenomena were notably absent, including processing of 3D shape, patterns on surfaces, occlusion, natural parts and a global advantage. Our results elucidate the computational challenges of vision by showing that learning to recognize objects suffices to produce some perceptual phenomena but not others and reveal the perceptual properties that could be incorporated into deep networks to improve their performance.

## INTRODUCTION

> How do I know this is true?
>
> I look inside myself and see.
>
> — Tao Te Ching (Mitchell, 1988)

Convolutional or deep neural networks have revolutionized computer vision with their human-like accuracy on vision tasks, and their object representations match coarsely with the brain (see Serre, 2019; Sinz et al., 2019 for detailed reviews). Yet, at a finer scale, they are still outperformed by humans (Katti et al., 2017; Katti and Arun, 2019) and show systematic deviations from human perception (Pramod and Arun, 2016a; Geirhos et al., 2018b; Rajalingham et al., 2018; Dodge and Karam, 2019). Even these differences are largely quantitative in that there are no explicit or emergent properties that are present in humans but absent in deep networks. This has given rise to the prevailing belief that any remaining differences between brains and deep networks can be fixed by training on larger datasets, incorporating more constraints (Sinz et al., 2019) or by making relatively minor modifications to network architecture such as by including recurrent feedback (Kar et al., 2019a; Kietzmann et al., 2019).

Despite these insights, we do not yet know whether there are qualitative differences between how brains and deep networks see. This is an important question because resolving qualitative differences might require non-trivial changes in network training or architecture. One approach could be to train deep networks on multiple visual tasks and compare them with humans, but the answer would be insightful only if networks fail to learn certain tasks (Fleuret et al., 2011a). Alternatively, we could compare qualitative or emergent properties of our perception with that of deep networks, provided these properties can indeed be checked in any deep network without explicit training for these properties.

Fortunately, many classic findings from visual psychology and neuroscience report emergent phenomena and properties that can be directly tested on deep networks. Consider for instance, the classic Thatcher effect (Figure 1A), in which a face with rotated parts looks grotesque in an upright orientation but looks entirely normal when inverted (Thompson, 1980). This effect can be recast as a statement about the underlying face representation: in perceptual space, the distance between the normal and Thatcherized face is presumably larger when they are upright than when they are inverted (Figure 1B). This has been confirmed using dissimilarity ratings in humans (Bartlett and Searcy, 1993). These distances can be compared for any representation, including for a deep network (Figure 1C). Since deep networks are organized layer-wise with increasing complexity across layers, this would also reveal the layers at which the deep network begins to experience or “see” a Thatcher effect (Figure 1D).

**Figure 1:**
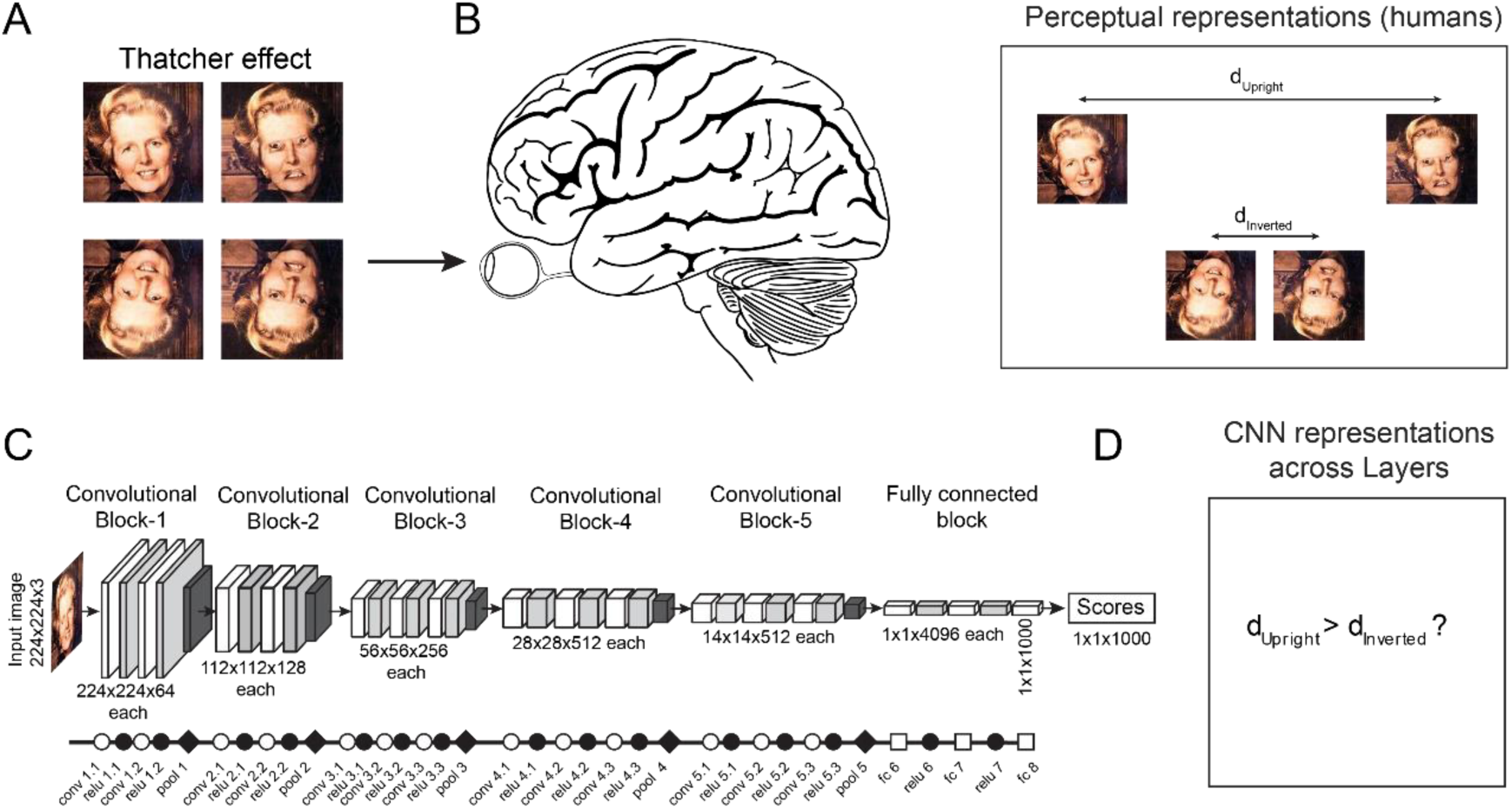
Evaluating whether deep networks see the way we do. (A) In the classic Thatcher effect, when the parts of a face are individually inverted, the face appears grotesque when upright (*top row*) but not when inverted (*bottom row*). Figure credit: Reproduced with permission from Peter Thompson. (B) When the brain views these images, it presumably extracts specific features from each face so as to give rise to this effect. We can use this idea to recast the Thatcher effect as a statement about the underlying perceptual space. The distance between the normal and Thatcherized face is larger when they are upright compared to when the faces are inverted. This property can easily be checked for any computational model. (C) Architecture of a common deep neural network (VGG-16). Symbols used here and in all subsequent figures indicate the underlying mathematical operations of that layer: *unfilled circle* for convolution, *filled circle* for ReLu, *diamond* for maxpooling and *unfilled square* for fully connected layers. Broadly, unfilled symbols depict linear operations and filled symbols depict non-linear operations. (D) By comparing the distance between upright and inverted Thatcherized images, we can ask whether any given layer of the deep network sees a Thatcher effect.

Knowing whether a deep network exhibits the Thatcher effect can be insightful for a variety of reasons. First, it would confirm that the deep network indeed does see faces the way we do. Second, this question can be asked of any deep network without explicit training to produce a Thatcher effect. For instance, testing this question on face and object detection networks would reveal whether object or face-specific training is sufficient for the emergence of the Thatcher effect. Finally, this question has relevance to neuroscience, because object representations in the early and late layers of deep networks match with early and late visual processing stages in the brain (Cichy et al., 2014; Kar et al., 2019b). The layer at which this effect arises could therefore reveal its underlying computational complexity and offer clues as to its neural substrates.

Here, we identified a number of emergent perceptual and neural properties from visual psychology and neuroscience that can be recast as statements about distances between images in the underlying perceptual/neural representation. We then tested each of these properties on a state-of-the-art deep neural network optimized for object recognition. This revealed a highly interesting and insightful list of properties that were either present or absent in the network.

## RESULTS

We identified a large number of emergent perceptual and neural properties that can be tested across layers of a deep network. We organized these properties broadly into five groups: (1) those that reveal sensitivity to object or scene statistics, comprising the Thatcher effect, mirror confusion and object-scene incongruence; (2) neural principles observed in high-level visual cortex, related to multiple object tuning and sparseness; (3) relational properties such as Weber’s law, relative size and surface invariant pattern processing; (4) encoding of 3d shape or scene structure and (5) processing of object parts and global structure.

We evaluated these properties for a state-of-the-art pre-trained deep network, VGG-16, optimized for object classification on the ImageNet dataset (Simonyan and Zisserman, 2014). For each property, we performed an experiment in which we used carefully controlled sets of images as input to the network, obtained the activations of the units in each layer, and asked whether each layer shows that property. We obtained qualitatively similar results on several other pre-trained feedforward networks with diverse architectures: AlexNet, GoogleNet, Resnet-50 and Resnet-152 (Section S1). To ensure that the results are truly due to training and not simply a consequence of the architecture of the network, we also repeated these experiments on a randomly initialized VGG-16 model (Section S2). For simplicity, we report our results only for the VGG-16 network in the sections below, with results for all other networks detailed in supplementary material (Sections S1-2).

### Experiment 1: Do deep networks see a Thatcher effect?

The Thatcher effect is an elegant demonstration of how upright faces are processed differently from inverted faces, presumably because we encounter mostly upright faces. As detailed in the Introduction, it can be recast as a statement about the underlying distances in perceptual space: that normal vs Thatcherized faces are closer when inverted than when upright (Figure 2A). For each layer of the deep network (VGG-16), we calculated a “Thatcher index” of the form (d_upright_ – d_inverted_)/(d_upright_ + d_inverted_), where d_upright_ is the distance between normal and Thatcherized face in the upright orientation, and d_inverted_ is the distance between them in an inverted orientation. Note that the Thatcher index for a pixel-like representation (where the activation of each unit is proportional to the brightness of each pixel in the image) will be zero since d_upright_ and d_inverted_ will be identical. For human perception, since d_upright_ > d_inverted_, the Thatcher index will be positive and can be estimated from previous studies (see Methods).

**Figure 2:**
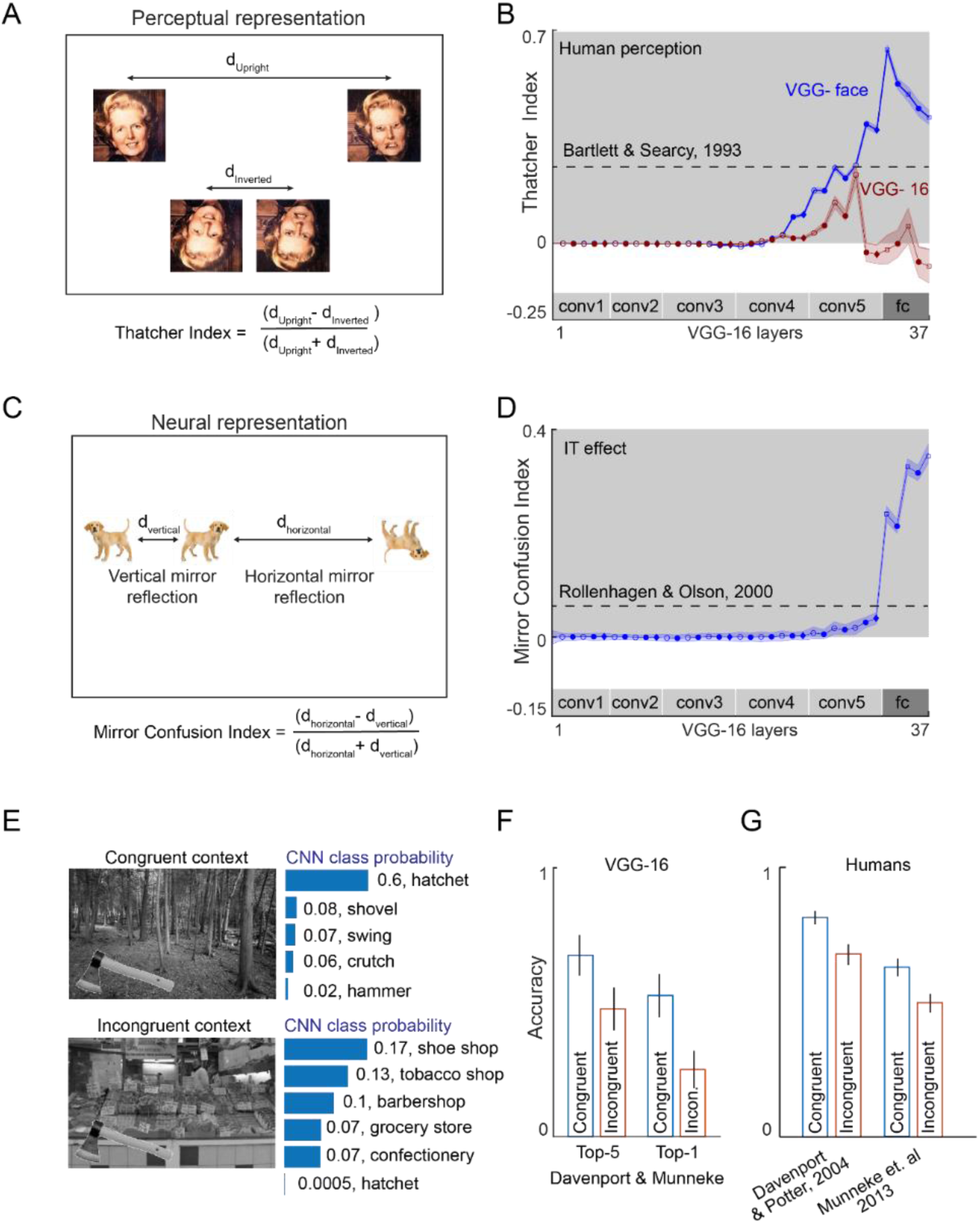
Object and scene regularities in deep networks. (A) Schematic showing the perceptual arrangement of normal and Thatcherized faces in the upright and inverted orientations. A Thatcherized face is more similar to its normal counterpart in the inverted orientation. (B) Thatcher index (averaged across 20 pairs of normal-Thatcherized face pairs) plotted across layers for two pre-trained networks, VGG-16 (red) and VGG-face (blue). Shaded error bars indicated s.e.m across 20 face pairs. The gray zone indicates human-like performance, with dotted lines indicating the strength of the Thatcher effect estimated from humans (see Methods). (C) Schematic showing the representation of vertical and horizontal mirror images in perception. Vertical mirror image pairs are closer than horizontal mirror image pairs. The same effect holds even if the dog is rotated by 90°, showing that this effect is not simply due to the object’s axis of elongation. (D) Mirror Confusion Index (averaged across 50 naturalistic objects and their 90° rotated versions) across layers for the VGG-16 network. Dotted lines show the strength of the mirror confusion index estimated from monkey inferior temporal neurons (see Methods). (E) An example object (*hatchet*) embedded in a congruent context (*forest*) and an incongruent context (*supermarket*), highlighting the vulnerability of deep network classification to scene context. The CNN class probability returned by the VGG-16 network is shown beside each image. It can be seen that the deep network returns the correct class label on the congruent but not the incongruent scene. (F) Accuracy of object classification by the VGG-16 network across congruent (blue) and incongruent(red) context scenes, for top-5 accuracy (*left*) and top-1 accuracy (*right*). Error bars represent s.e.m across 40 congruent/incongruent scene pairs. (G) Accuracy of object naming by humans in congruent (blue) and incongruent (red) contexts across two separate studies (Davenport and Potter, 2004; Munneke et al., 2013). Humans are less accurate on naming objects placed on incongruent scene contexts, but show a smaller drop compared to the VGG-16 network. Error bars represent s.e.m reproduced from the published studies.

Next, we calculated the Thatcher index across layers for two networks with similar architecture but trained on different tasks (see Methods). The first was VGG-16 which is trained for object classification (Simonyan and Zisserman, 2014). The second was VGG-face which is trained for face recognition (Parkhi et al., 2015). The Thatcher index for both networks across layers is shown in Figure 2B. It can be seen that the VGG-16 shows a positive Thatcher index in the conv4 and conv5 layers but eventually does not show a Thatcher effect in the final fully connected layers (Figure 2B, red curve). By contrast, the VGG-face network shows a steadily rising Thatcher effect that remains close to human levels even in the fully connected layers. Thus, the Thatcher effect is present only in deep networks trained on upright face recognition but not on generic object recognition.

### Experiment 2: Do deep networks show mirror confusion?

Mirror reflections along the vertical axis appear more similar to us than reflections along the horizontal axis (Figure 2C). This effect has been observed both in behaviour as well as in high-level visual cortex in monkeys (Rollenhagen and Olson, 2000). To assess whether deep networks show mirror confusion, we calculated a mirror confusion index of the form (d_horizontal_ – d_vertical_)/(d_horizontal_ + d_vertical_), where d_horizontal_ and d_vertical_ represent the distance between horizontal mirror image pairs and between vertical mirror image pairs respectively (see Methods). Since vertical mirror images are more similar in perception, this index would be positive. Across the deep network VGG-16, we found an increasing mirror confusion index across layers (Figure 2D). Thus, the deep network does experience (as we do) more mirror confusion for vertical compared to horizontal mirror images.

### Experiment 3: Do deep networks show scene incongruence?

Our ability to recognize an object is hampered when it is placed in an incongruent context (Davenport and Potter, 2004; Munneke et al., 2013), suggesting that our perception is sensitive to the statistical regularities of objects co-occurring in specific scene context. To explore whether deep networks are also sensitive to scene context, we gave as input the same images as tested on humans, and asked how the deep network classification output changes with scene context (see Methods for more details). An example object (hatchet) placed against a congruent context (forest) and incongruent context (supermarket) are shown in Figure 2E. The VGG-16 network returned a high probability score for the correct target object in the congruent context (Figure 2E, top row) but gave a low probability score for the same object in an incongruent context (Figure 2E bottom row). We obtained similar results across all congruent/incongruent scene pairs: the VGG-16 top-1 accuracy dropped substantially for incongruent compared to congruent contexts (drop in accuracy from congruent to incongruent scenes: 20% for top-5 accuracy; 27% for top-1 accuracy; Figure 2F). On the same scenes, human object naming accuracy has been reported to drop for incongruent scenes, but the drop was smaller compared to the VGG-16 network (drop in human accuracy from congruent to incongruent scenes: 14% in the Davenport & Potter, 2004; 13% in Munneke et. al. 2013; Figure 2G). We note that assessing the statistical significance of the accuracy difference in humans and deep networks is not straightforward since the variations in accuracy reported are across subjects for humans and across scenes for the VGG-16 network. We conclude that deep networks show scene incongruence albeit to a smaller extent than humans.

### Experiment 4: Do deep networks show multiple object normalization?

Next we asked whether individual units in deep networks conform to two general principles observed in single neurons of high-level visual cortex. The first one is multiple object normalization, whereby the neural response to multiple objects is the average of the individual object responses at those locations (Zoccolan et al., 2005). This principle is illustrated in Figure 3A. Note that this analysis is meaningful only for units that respond to all three locations: a unit in an early layer with a small receptive field would respond to objects at only one location regardless of how many other objects were present in the image. To identify units that are responsive to objects at each location, we calculated the variance of its activation across all objects presented at that location. We then performed this analysis on units that showed a non-zero response variance at all three locations, which meant units in Layer 23 (conv4.3) onwards.

**Figure 3:**
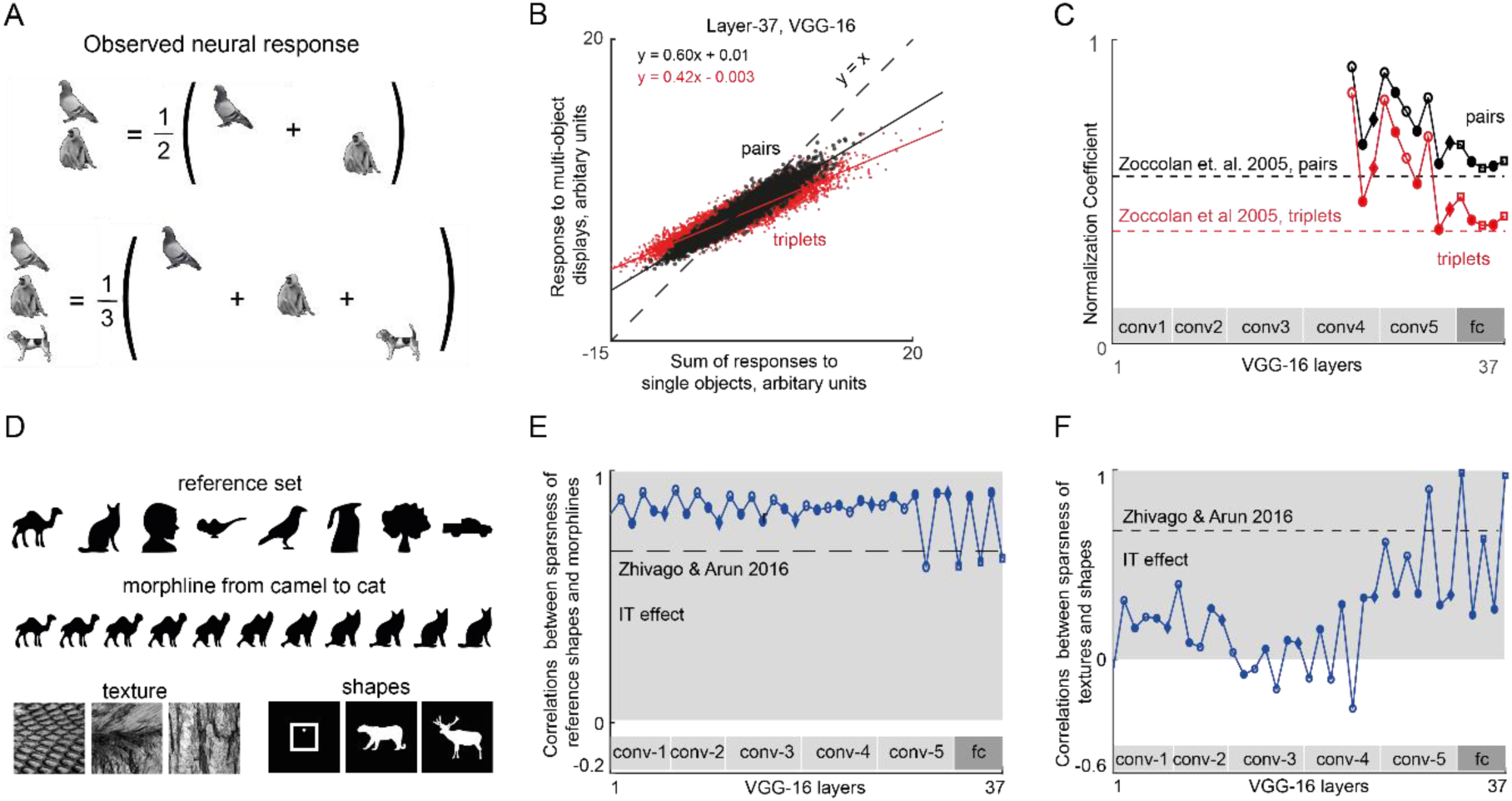
Single unit properties of deep networks. (A) Schematic illustrating a general principle observed in neurons in high-level visual areas of the brain. The response of a neuron to multiple objects is typically the average of its responses to the individual objects at those locations. (B) Response to multiple object displays plotted against the sum of the individual object responses for two-object displays (black) and three-object displays (red), across 10,000 units randomly selected from Layer 37 of the VGG-16 network. (C) Normalization slope plotted across layers for two object displays (black) and three-object displays (red). (D) Stimuli used to compare selectivity across multiple dimensions in a previous study of IT neurons (Zhivago and Arun, 2016). (E) Correlation between sparseness on the reference set and maximum sparseness along morphlines across units of each layer in the VGG-16 network. (F) Correlation between sparseness along textures and sparseness along shapes plotted across layers of the VGG-16 network.

To assess whether deep networks show multiple object normalization, we plotted for each unit in a given layer its response to multiple objects against the sum of its responses to the individual objects. If there is multiple object normalization, the slope of the resulting plot should be 1/2 for two objects and 1/3 for three objects. The resulting plot is shown for Layer 37 of the VGG-16 network (Figure 3B). The overall slope was 0.60 for two objects and 0.42 for three objects for all units. To evaluate this effect across layers, we plotted the two-object and three-object slopes obtained in this manner across layers (Figure 3C). For the later layers we observed a nearly monotonic decrease in the slopes, approaching the levels observed in monkey high-level visual areas (Figure 3C). We conclude that deep networks exhibit multiple object normalization.

### Experiment 5: Do deep networks show selectivity across multiple dimensions?

In a recent study we showed that neurons in the monkey inferior temporal cortex have intrinsic constraints on their selectivity that manifests in two ways (Zhivago and Arun, 2016). First, neurons that respond to fewer shapes have sharper tuning to parametric changes in these shapes. To assess whether units in the deep network VGG-16 show this pattern, we calculated the sparseness of each unit across a reference set of disparate shapes (Figure 3D), and its sparseness for parametric changes between pairs of these stimuli (an example morph line is shown in Figure 3D). This revealed a consistently high correlation across units of each layer in the VGG-16 network (Figure 3E). Second, we found that neurons that are sharply tuned across textures are also sharply tuned to shapes. To assess this effect across layers, we calculated the correlation across units between sparseness on textures with the sparseness on shapes. Although there was no such consistently positive correlation in the early layers, we did find a positive correlation in the later (conv5 & fc) layers (Figure 3F). We conclude that deep networks show selectivity along multiple dimensions just like neurons in high-level visual cortex.

### Experiment 6: Do deep networks show Weber’s law?

Next we asked whether deep networks are sensitive to relational properties in visual displays. The first and most widely known of these is Weber’s law, which states that sensitivity to changes in any sensory quantity is proportional to the baseline level being used for comparison. Weber’s law for line length is illustrated in Figure 4A. This in turn predicts that the distance between any two lines differing in length should be proportional to the relative but not absolute change in length. In a previous study, we showed that this is true for humans in visual search for both length and intensity changes (Pramod and Arun, 2014).

**Figure 4:**
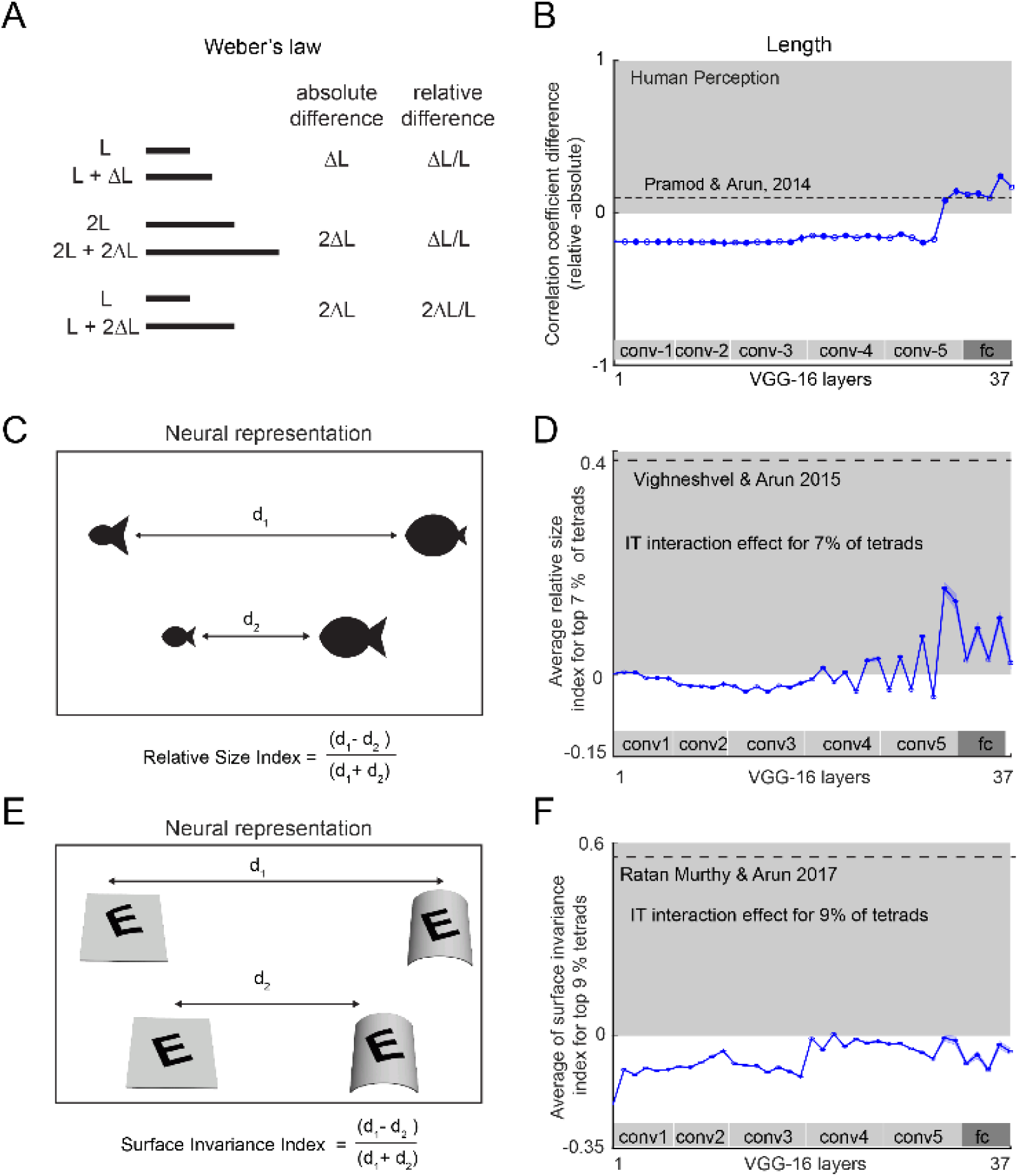
Relational properties in deep networks. (A) Example illustrating the Weber’s law for line length. Although the original statement of Weber’s law is that the just-noticeable difference in length will depend on the baseline length, it also applies to perceptual distances between any two stimuli (Pramod and Arun, 2014). In this formulation, the perceptual distance between two lines differing in length will be proportional to the relative change in length rather than the absolute change. (B) To calculate a single quantity that measures adherence to Weber’s law, we calculated the difference in correlation between pairwise distances between lines with the relative and absolute differences in line length. A positive difference indicates adherence to Weber’s law (indicated by the gray shading). This difference in correlation is plotted across layers for line length. (C) Schematic of the relative size encoding observed in monkey IT neurons (Vighneshvel and Arun, 2015). For a fraction of neurons, the distance between two-part objects when both parts covary in size is smaller than the distance when they show inconsistent changes in size. Thus, these neurons are sensitive to the relative size of items in a display. (D) Relative size index across units with interaction effects (averaged across top 7% tetrads tetrads, error bars representing s.e.m) across layers of the VGG-16 network. Dotted lines show the strength of the relative index estimated from monkey inferior temporal neurons (Vighneshvel and Arun, 2015). (E) Schematic of the surface invariance index observed in monkey IT neurons (Ratan Murty and Arun, 2017). For a fraction of neurons, the distance between two stimuli with congruent changes in pattern and surface curvature is smaller than between two stimuli with incongruent pattern/surface changes. Thus, these neurons decouple pattern shape from surface shape. (F) Surface invariance index across units with interaction effects (averaged across top 9% pattern/surface tetrads, error bars representing s.e.m) across layers of the VGG-16 network. Dotted lines show the strength of the surface invaraince index estimated from monkey inferior temporal neurons (Ratan Murty and Arun, 2017).

We therefore asked for the deep network VGG-16, whether pairwise distances between lines of varying length are correlated with absolute or relative changes in length. Specifically, if the correlation between pairwise distances and relative changes in length is larger than the correlation with absolute changes in length, we deemed that layer to exhibit Weber’s law. This difference in correlation is positive for humans in visual search, and we plotted this difference across layers of the VGG-16 network (Figure 4B). The correlation difference was initially negative in the early layers of the network, meaning that the early layers were more sensitive to absolute changes in length. To our surprise, however, distances in the later layers were sensitive to relative changes in length in accordance to Weber’s law (Figure 4B).

We conclude that deep networks exhibit Weber’s law for length.

### Experiment 7: Do deep networks encode relative size?

We have previously shown that neurons in high-level visual areas are sensitive to the relative size of items in a display (Vighneshvel and Arun, 2015). Specifically, we found that, when two items in a display undergo congruent changes in size, the neural response is more similar than expected given the two individual changes in size. This pattern was present only in a small fraction (7%) of the neurons. This effect is illustrated in Figure 4C. To explore whether this effect can be observed in a given layer of the deep network VGG-16, we performed a similar analysis. We selected units in a given layer with the strongest interactions (see Methods) and calculated a relative size index of the form (d1-d2)/(d1+d2) where d1 is the distance between stimuli with incongruent changes in size, and d2 is the distance between stimuli with congruent size changes (i.e. where both items or parts are scaled up or down in size by the same factor). The relative size index was calculated for each tetrad as to replicate the same analysis done in the previous study(Vighneshvel and Arun, 2015). The relative size index for the VGG-16 network remained close to zero in the initial layers and increased modestly to a positive level in the later layers (Figure 4D). However the size of this effect was far smaller than that observed in IT neurons, but in the same direction. We conclude that the deep network VGG-16 encodes relative size.

### Experiment 8: Do deep networks decouple pattern shape from surface shape?

A recent study showed that IT neurons respond more similarly when a pattern and a surface undergo congruent changes in curvature or tilt (Ratan Murty and Arun, 2017). This effect is illustrated for a pattern surface pair in Figure 4E, where it can be seen that the distance between incongruent pattern-surface pairs (where the pattern and surface change in opposite directions) is larger than the distance between congruent pairs where the pattern and surface undergo congruent changes. To assess whether the deep network VGG-16 shows this property, we identified units with increased interactions (see Methods) and calculated a surface invariance index of the form (d1-d2)/(d1+d2), where d1 is the distance between incongruent pairs, and d2 is the distance between congruent pairs. A positive value of this index for a given layer implies that the layer shows surface invariance. However the surface invariance index was consistently below zero across layers for the VGG-16 network (Figure 4F). We conclude that the VGG-16 network does not show surface invariance.

### Experiment 9: Do deep networks show 3d processing?

We are sensitive to three-dimensional shape and not simply two-dimensional contours in the image. This was demonstrated in an elegant experiment in which search for a target differing in 3d structure is easy whereas search for a target with the same difference in 2d features is hard (Enns and Rensink, 1990, 1991). This effect can be recast as a statement about distances in perceptual space as illustrated in Figure 5A. All three pairs of shapes depicted in Figure 5A differ in the same Y-shaped feature, but the two cuboids are more dissimilar because they differ also in 3d shape. To assess whether units in a given layer of the deep network show this effect, we calculated a 3d processing index of the form (d1-d2)/(d1+d2) where d1 is the distance between the cuboids and d2 is the distance between the two equivalent 2d conditions. A positive 3D processing index indicates an effect similar to human perception. However we found that the 3D processing index was consistently near zero or negative across all layers of the VGG-16 network (Figure 5B). We conclude that deep networks are not sensitive to 3d shape. We speculate that explicit training of deep networks on 3d shape processing may be required for the network to exhibit this effect.

**Figure 5:**
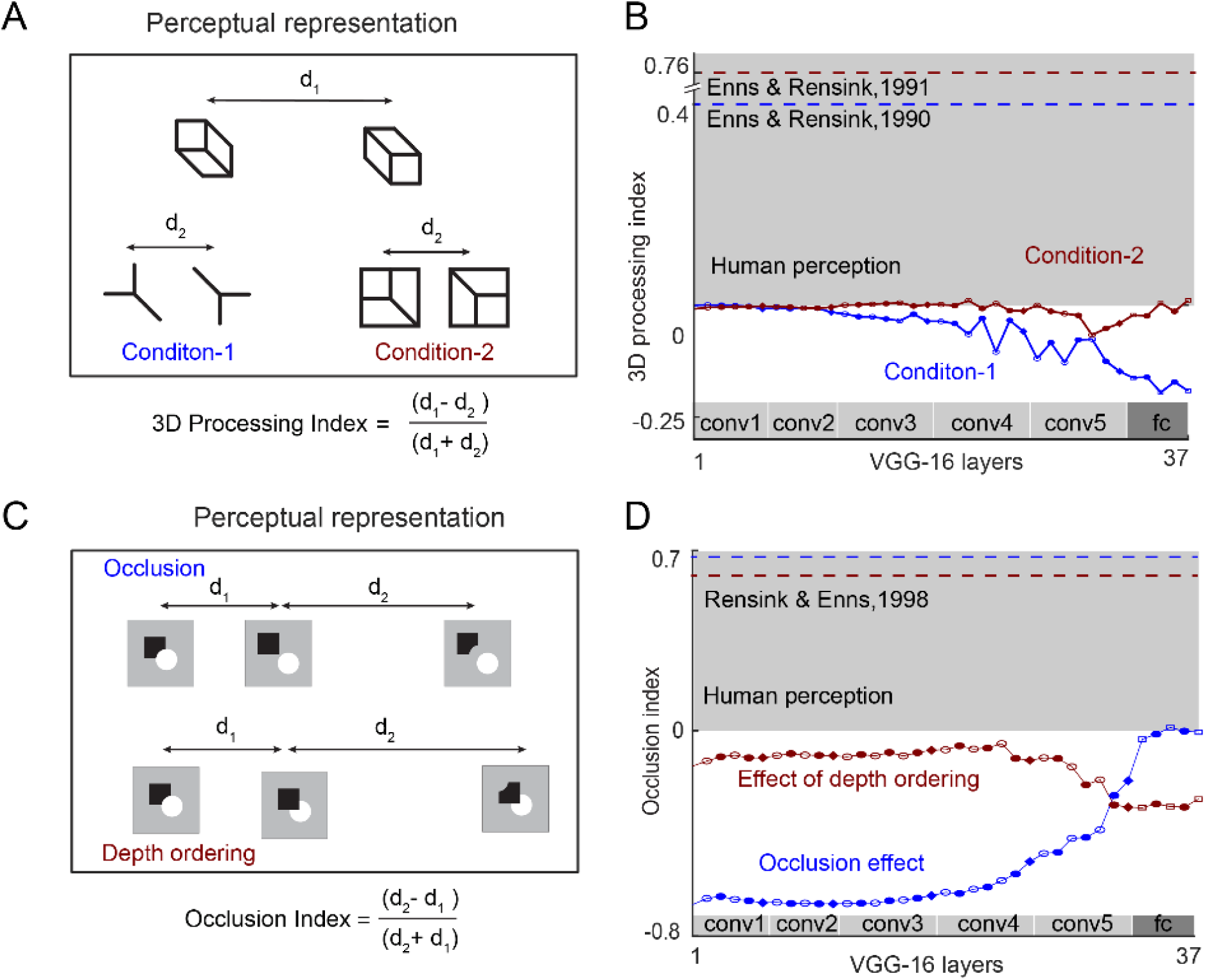
3D processing in deep networks. (A) Schematic demonstrating sensitivity to 3d shape in our perception (Enns and Rensink, 1991). Three equivalent image pairs are shown in perceptual space. The first image pair (with distance marked d1) consists of two cuboids at different orientations, and corresponds to an easy visual search i.e. the two objects are less similar. The second pair (marked with distance d2) contains the same feature difference as in the first pair, but represents a hard search i.e. is perceived as much more similar. Likewise, the third pair, with the same feature difference as the other two, is a hard search i.e. perceived as similar. (B) 3D processing index for the VGG-16 network across layers, for condition 1 (d_1_ vs d_2_, *blue*) and condition 2 (d_1_ vs d_3_, *red*). Dotted lines of the corresponding color represent the estimated human effect size. (C) Schematic showing processing of occlusions in perception*. Top:* A square alongside a disk is perceptually similar to a display with the same objects but occluded, but dissimilar to a 2d control image with an equivalent feature difference. *Bottom:* A square occluding a disk or disk occluding square are perceptually similar, but dissimilar to an equivalent 2d control with the same set of feature differences. (D) Occlusion index for both basic occlusion (*blue*) and depth ordering (*red*) for each layer of the VGG-16 network.

### Experiment 10: Do deep networks understand occlusions?

A classic finding in human perception is that we automatically process occlusion relations between objects (Rensink and Enns, 1998). Specifically, search for a target containing occluded objects among distractors that contain the same objects unoccluded is hard, whereas searching for the equivalent 2d feature difference is much easier (Figure 5C, top row). Likewise, searching for a target that is different in the order of occlusion is hard, whereas searching for the equivalent 2d feature difference is easy (Figure 5C, bottom row). These observations demonstrate that our visual system has a similar representation for occluded and unoccluded objects.

We therefore asked whether similar effects could be observed in the VGG-16 network, by calculating an occlusion index of the form (d2-d1)/(d2+d1) where d1 is the distance between two displays that are equivalent except for occlusion, and d2 is the distance between equivalent displays with the same 2d feature difference. A positive occlusion index implies an effect similar to human perception. The occlusion index remained consistently below across all layers of the VGG-16 network, for both the occlusion effect and occlusion ordering (Figure 5D). We conclude that deep networks do not understand occlusions.

### Experiment 11: Do deep networks understand object parts?

We not only recognize objects but are able to easily describe their parts. We conducted two related experiments to investigate part processing in deep networks. In Experiment 11A, we compared deep network feature representations for whole objects and for the same object broken down into either natural or unnatural parts. In perception, searching for an object broken into its natural parts with the original object as distractors is much harder than searching for the same object broken at an unnatural location (Xu and Singh, 2002). This result is depicted schematically in terms of the underlying distances (Figure 6A). To assess whether the VGG-16 network also shows this part decomposition, we calculated a part processing index of the form (d_u_ – d_n_)/(d_u_ + d_n_) where d_u_ is the distance between the original object and the object broken at an unnatural location, and d_n_ is the distance between the original object and the same object broken at a natural location. A positive part processing index implies an effect similar to that seen in perception. The part processing index across layers of the VGG-16 network is depicted in Figure 6B. We found that the index becomes positive in the intermediate layers, but becomes negative in the subsequent layers (conv4/conv5 onwards).

**Figure 6:**
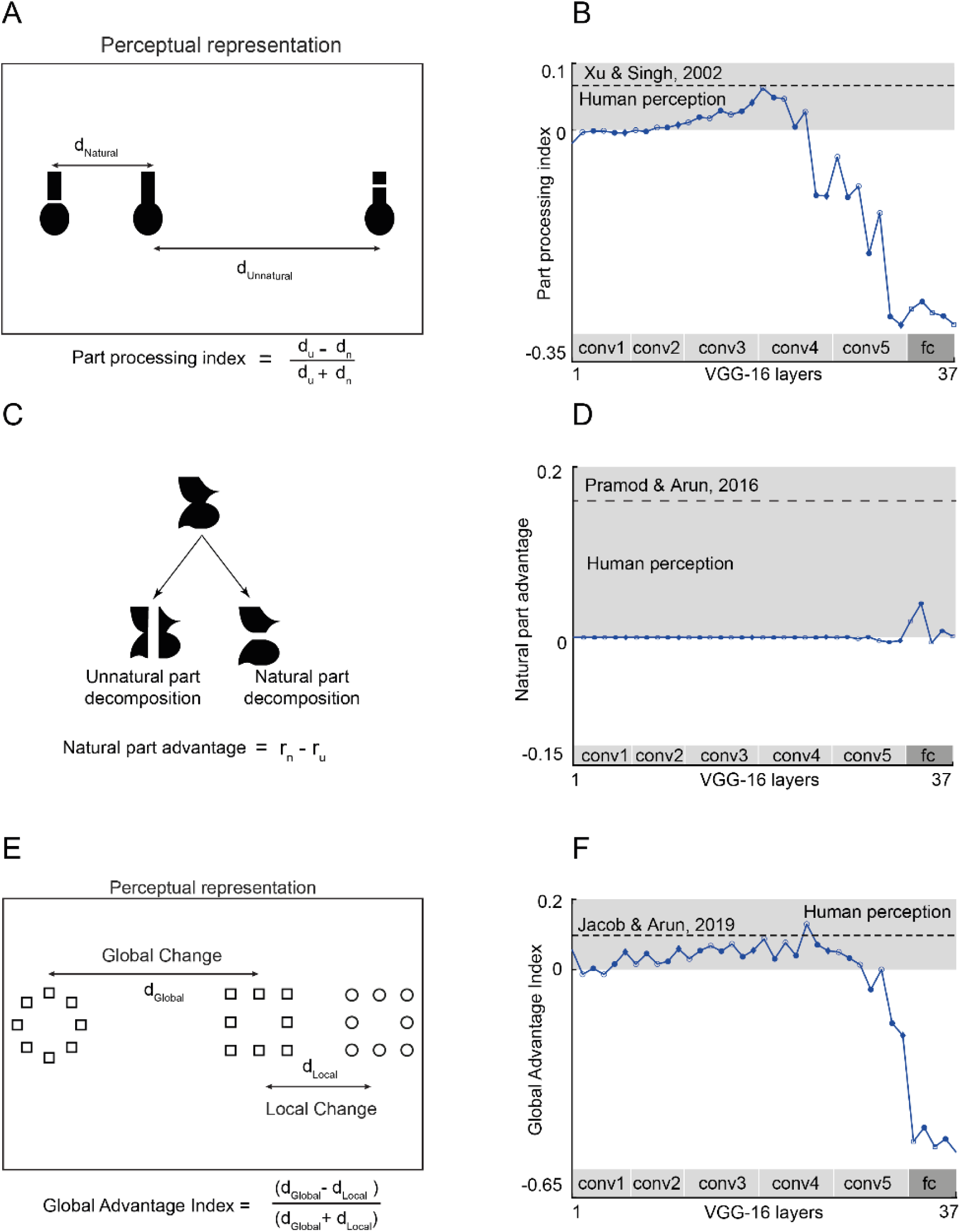
Part-whole relations in deep networks. (A) Schematic showing the perceptual representation of objects with a break introduced either at natural or unnatural parts. (B) Part processing index across layers of the VGG-16 network. The shaded gray bar represents effects similar to human perception, and the dotted line represents the effect size estimated from human visual search (Xu and Singh, 2002). (C) Schematic showing how the same object can be broken into either natural or unnatural parts. (D) Natural part advantage across layers of the VGG-16 network. The shaded gray bar represents effects similar to human perception, and the dotted line represents the effect size estimated from human visual search (Pramod and Arun, 2016b). (E) Schematic showing the perceptual representation of hierarchical stimuli. The left and middle images differ only in global shape whereas the middle and right images differ only at the local level. (F) Global Advantage index across layers of the VGG-16 network. The shaded gray bar represents effects similar to human perception, and the dotted line represents the effect size estimated from human visual search (Jacob and Arun, 2019).

In Experiment 11B, we asked what happens to objects that can be decomposed into two possible ways without introducing a break (Figure 6C). In visual search, search between pairs of whole objects is better explained by breaking the object down into its natural parts compared to its unnatural parts (Pramod and Arun, 2016b). To capture this effect, we defined the natural part advantage as the difference in model correlation (see Methods) between natural and unnatural parts. A positive value indicates effects similar to perception. This natural part advantage is shown across layers of the VGG-16 network in Figure 6D. It can be seen that there is little or no advantage for natural parts in most layers except temporarily in the later layers (conv5/fc).

Based on Experiments 11A & 11B, we conclude that the VGG-16 network shows no systematic part processing.

### Experiment 12: Do deep networks show a global shape advantage?

In perception a classic finding is that we see the forest before the trees, i.e. we can detect global shape before local shape (Navon, 1977; Kimchi, 1994). We can recast this effect into a statement about distances in perception: the distance between two hierarchical stimuli differing only in global shape will be larger than the distance between two such stimuli differing only in local shape. This is depicted schematically in Figure 6E. To calculate a single measure for this effect, we defined a global advantage index as (d_global_ – d_local_)/(d_global_ + d_local_), where d_global_ is the average distance between all image pairs differing only in global shape and d_local_ is the average distance between all image pairs differing only in local shape. A positive global advantage index implies an effect similar to perception. The global advantage index is depicted across layers of the VGG-16 network in Figure 6F. While there is a slight global advantage in the initial layers, the network representation swings rapidly in the later layers towards the opposite extreme, which is a local advantage. Thus, deep networks see the trees and not the forest.

## DISCUSSION

Because object representations in deep networks match coarsely with the brain, it is widely believed that any remaining differences between brains and deep networks must be of degree but not of kind. Here, we show that this is not always the case: In some respects, deep networks do see the way we do. They exhibit perceptual phenomena such as the Thatcher effect, mirror confusion, scene incongruence and Weber’s law. Their units show multiple object normalization, sparseness along multiple dimensions and encode relative size. Yet in other ways, they don’t see the way we do. They fail to encode relational properties such as surface invariance, do not show processing of 3d features, occlusions or natural parts and do not show a global advantage.

These findings are important for several reasons. First, they describe the similarities and differences between our vision and deep networks, and challenge the prevailing belief that they can be treated as accurate models for biological vision. Second, they show that object recognition training alone is sufficient to produce some emergent properties but not others, thereby elucidating the computational problem of vision itself. Finally, the missing properties could be incorporated as training or architecture constraints on deep networks to yield better or more robust performance. Below we discuss our findings in the context of the existing literature.

We begin by discussing some general concerns regarding our findings. First and foremost, it could be argued that our results are based on testing with artificial objects or images, and that it is unreasonable to expect deep networks to respond sensibly to unnatural images. However, these concerns apply equally to humans as well, who in fact do respond sensibly to these artificial displays despite no prior exposure. Indeed, there is a long tradition in psychology and neuroscience of using artificial images to elucidate visual processing (Rust and Movshon, 2005). Second, it could be argued that deep networks could potentially be trained to report all the tested properties. However, such a finding would only be circular if the network did indeed exhibit the property it was trained for. We do note however that it would be interesting if deep networks were unable to learn certain properties. Indeed, certain relational properties have been reported as difficult to learn by computer vision algorithms (Fleuret et al., 2011b), although this study did not evaluate deep neural networks. By contrast, we consider our experiments to be far more interesting, since they reveal that deep networks trained for specific other purposes show emergent properties that they were not explicitly trained for, such as Weber’s law.

Our finding that deep networks trained for object recognition exhibit Weber’s law or encode relative size is puzzling at first glance. Why would the demands of recognizing objects require sensitivity to relative changes? One possibility is that object recognition requires a representation invariant to changes in size, position, viewpoint and even illumination of objects in the image, which in turn requires processing all object features relative to the surrounding features. This could be tested by training deep networks on controlled sets of images with variations of one kind but not the other. We note that there could be other visual task requirements that could also give rise to Weber’s law (Pardo-Vazquez et al., 2019).

Our finding that deep networks exhibit the Thatcher effect, mirror confusion and scene incongruence are consistent with them being sensitive to image regularities in scenes. In fact, the VGG-16 network may be over-reliant on scene context, because it showed a larger drop in accuracy for incongruent scenes compared to humans (Figure 2F). This is consistent with a previous study in which human scene expectations benefited deep network performance (Katti et al., 2019). However, the VGG-16 network did not exhibit 3d processing, occlusions and surface invariance, suggesting that these properties might emerge only with additional task demands such as extracting 3d shape from the image. Likewise, the absence of any part processing or global advantage in the network suggests that these too are properties that might emerge with additional task demands, such as part recognition or global form recognition (Belongie et al., 2002).

We have found that deep networks do not show a global advantage effect but instead seem to process local features. This finding is surprising considering that units in later layers receive convergent inputs from the entire image. However, our finding is consistent with reports of a bias towards local object texture in deep networks (Geirhos et al., 2018a). It is also consistent with the large perturbations in classification observed when new objects are added to a scene (Rosenfeld et al., 2018), which presumably change the distribution of local features. Our finding that deep networks experience large scene incongruence effects is therefore likely to be due to mismatched local features rather than global features. Indeed, incorporating scene expectations from humans (presumably driven by global features) can lead to substantial improvements in object recognition (Katti et al., 2019). Finally, a reliance on processing local features is probably what makes deep networks detect incongruously large objects in scenes better than humans (Eckstein et al., 2017). We speculate that training on global shape could make deep networks more robust and human-like in their performance.

Comparing human vision and deep neural networks depends critically on the choice of network architecture, learning algorithm, dataset and the task learned. An important but neglected finding is that even a randomly initialized network can generate features useful for certain tasks. For tasks like texture generation and discrimination, even a randomly initialized network can yield reliable features (Jarrett et al., 2009; Mongia et al., 2016). Likewise, a significant amount of explainable variance in the early visual responses of MEG signals in humans can be predicted using features extracted from a randomly initialized and untrained deep neural network (Cichy et al., 2016). By contrast, we have found that most perceptual effects are absent in randomly initialized networks, except for global advantage (Section S2). Thus, training on object classification abolishes the global advantage and introduces sensitivity to local features.

How can deep networks be made to match neural and perceptual representations? There could be several ways of doing so. The first and perhaps most promising direction would be to explicitly train deep networks to produce such properties in addition to categorisation (Ruder, 2017). Another alternative would be to train deep networks on tasks such as navigation or agent-object interaction rather than (or in addition to) object recognition as this is ostensibly what humans also do (Haber et al., 2018; Yang et al., 2019).

Finally, we note that deep networks are notorious for their susceptibility to adversarial attacks. State-of-the-art deep neural networks have been shown to fail catastrophically when input images are subjected to carefully constructed changes that are barely perceivable to human eyes (Szegedy et al., 2013; Su et al., 2019). Likewise, deep networks can give erroneous predictions on completely nonsensical images (Nguyen et al., 2015). Finally, realistic multi-part 3D objects are consistently misclassified by deep networks across viewpoint changes (Athalye and Carlini, 2018). What could underlie such unusual behaviour? One possible reason could be the tendency for deep networks to prioritize local features as described earlier. We speculate that training deep networks to exhibit all the perceptual and neural properties described in this study might not only improve their performance but also make them more robust to adversarial attacks.

## METHODS

### VGG-16 network architecture & training

All experiments were performed on the VGG-16 network, a feedforward pre-trained deep convolutional neural network trained for object classification on the ImageNet dataset(Deng et al., 2009). We briefly describe the network architecture here, and the reader is referred to the original paper for more details (Simonyan and Zisserman, 2014). The input to the network is an RGB image of size 224 × 224 × 3 and the final output is a vector of confidence scores across 1000 categories. We subtracted the mean RGB value across all images (mean values across all images: R=123.68, G=116.78, B=103.94). The image is passed through a stack of convolutional filters (Figure 1C), where the initial layers have small receptive field (3×3) and later layers are fully connected. A non-linear rectification (ReLu) operation is done after each convolution operation. Five max-pooling layers are present to spatial pool the maximum signal over a 2−2 window of neurons. We used the MATLAB-based MatConvNet software platform (Vedaldi and Lenc, 2014) to extract features and do the analysis. In addition to VGG-16, we also used VGG-face which has the same architecture but trained instead on face identification (Parkhi et al., 2015).

### Feature Extraction

For each image, we passed it as input into the network, stored the activations of each layer as a column vector. Hence, a single image we will have 37 feature vectors (one column vector from each layer). To calculate the distance between images A and B, we calculated the Euclidean distance between the corresponding activation vectors.

In all the experiments, we define specific measure based on distances and this quantity is plotted across layers with a specific chain of symbols as shown in Figure 1C. Symbols used indicates the underlying mathematical operations done in that layer: unfilled circle for convolution, filled circle for ReLu, diamond for maxpooling and unfilled square for fully connected layers. Broadly, filled symbols denote linear operations and unfilled ones indicate non-linear operations.

### Experiment 1: Thatcher effect

The stimuli comprised 20 co-registered Indian faces (19 male, 1 female) from the IISc Indian face dataset (Katti and Arun, 2019). All faces were grayscale, upright and front-facing. To Thatcherize a face, we inverted the eyes and mouth while keeping rest of the face intact. We implemented inversion by first registering facial landmarks on frontal faces using an Active appearance model-based algorithm (Cootes et al., 2001). Briefly, this method models face appearance as a two-dimensional mesh with 76 nodes, each node represents local visual properties of stereotyped locations such as corners of eyes, nose, and mouth. We then defined bounding boxes for left and right eye as well as mouth, by identifying landmarks that correspond to the four corners of each box. We then locally inverted eye and mouth shape by replacing the top row of eye or mouth image pixels by the last row and likewise repeating this procedure for each pair of equidistant pixels rows above and below the middle of the local region. The inversion procedure was implemented as a custom script written in MATLAB. The full stimulus set is shown in Section S3.

To calculate a single measure for the Thatcher effect, we calculated a Thatcher index defined as 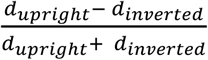, where *d_upright_* is the distance between an normal face and Thatcherized face in upright orientation and *d_inverted_* is the distance between a normal face and Thatcherized face in inverted orientation. We estimated the Thatcher index for humans from the similarity ratings reported from humans albeit on a different set of faces (Bartlett and Searcy, 1993). We calculated Thatcher index after converting the similarity rating (humans gave a rating between 1 to 7 on pair of images) into a dissimilarity rating (dissimilarity rating = 7-similarity rating).

### Experiment 2: Mirror Confusion

The stimuli consisted of 100 objects (50 naturalistic objects and 50 versions of these objects made by rotating each one by 90°). This was done to avoid any effect due to the objects own axis of elongation. We created a horizontal and vertical mirror image of each object. We then gave as input the original image and the two mirror images to the VGG-16 network and calculated for each layer the distance between the object and two mirror images. The full stimulus set is shown in Section S3.

To calculate a single measure for mirror confusion, we defined a mirror confusion index of the form 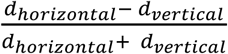, where *d_horizontal_* is the distance between an object and its horizontal mirror image and *d_vertical_* is the distance between an object and its vertical mirror image. We estimated the strength of mirror confusion index in the brain using previously published data from monkey IT neurons (Rollenhagen and Olson, 2000). Specifically, we took *d_horizontal_* to be the reported average firing rate difference between the original objects and its horizontal mirror image, and analogously for *d_vertical_*.

### Experiment 3: Scene Incongruence

The stimuli consisted of 40 objects which was taken from previous studies: 17 objects were from the Davenport study (Davenport and Potter, 2004) and the remaining 33 from the Munneke study (Munneke et al., 2013). We discarded a few objects from each set since they did not have a matching category in the ImageNet database. Each object was embedded against a congruent and an incongruent background. The full stimulus set is shown in Section S3.

We measured the classification accuracy (Top-1 and Top-5) of the VGG-16 network for the objects pasted onto congruent and incongruent scenes. The final layer of VGG-16 (Layer 38) returns a probability score for all 1000 categories in the ImageNet database. The top-1 accuracy is calculated as the average accuracy with which the object class with the highest probability matches the ground-truth object label. The top-5 accuracy is calculated as the average accuracy with which the ground-truth object is present among the object classes with the top 5 probability values. We report the human (object naming) accuracy on the same dataset from previous studies (Davenport and Potter, 2004; Munneke et al., 2013).

### Experiment 4: Multiple object normalization

The stimuli consisted of forty-nine natural grayscale images and placed them at three different locations in the image (Figure 4A). We have 147 (49 × 3 = 147) singletons and randomly selected 200 pairs and 200 triplet composites. We extracted features for all images (singletons, pairs and triplets) from every layer of the CNN. We selected a unit for further analysis only if the unit responded differently to at least one of the images in all the three positions. We then plotted the sum of activations of selected units in a layer to the singleton images against the activation for the corresponding pairs (or triplets). The slope of this scatterplot across layers was used to infer the nature of normalization in CNNs – a slope of 0.5 for pairs and 0.33 for triplets indicated divisive normalization matching that observed in high-level visual cortex. The full stimulus set is shown in Section S3.

### Experiment 5: Selectivity along multiple dimensions

Here we used the stimuli used in a previous study to assess the selectivity of IT neurons along multiple dimensions (Zhivago & Arun, 2016). These stimuli consisted of 8 reference shapes (Figure 3D; top row) and created intermediate parametric morphs between pairs of these shapes (Figure 3D; example morph between camel to cat). In addition, to compare texture and shape selectivity, we used 128 natural textures and 128 silhouette shapes. The full stimulus set is shown in Section S3.

As before we calculated the activation of every layer of the VGG-16 network to each of the above stimuli are input. Visually active neurons were selected by finding units with a non-zero variance across this stimulus set. We found the visually active neurons for each set separately and we selected a unit for further analysis only if that unit is visually active for both sets. The response of each unit was normalized between 0 and 1. We then calculated the sparseness of each unit across different stimulus sets: the reference set, the four morphlines, shape set and texture set. For a given stimulus set with responses r_1_, r_2_, r_3_, … r_n,_ where n is the number of stimuli, the sparseness is defined as follows: 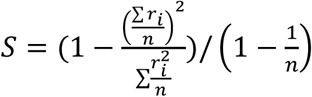 (Vinje et al., 2000; Zhivago and Arun, 2016). We then calculated the correlation across neurons between the sparseness on one stimulus set versus another stimulus set.

### Experiment 6: Weber’s Law

To test for the presence of Weber’s law in the deep network, we manipulated two elementary features: length and intensity. We created two sets of images, one varying in the length and the other varying in brightness of a rectangular bar. We chose the values of lengths and brightness’s such that the feature difference computed on pairs of images spanned a wide range both in terms of absolute as well as relative differences. The full stimulus set is shown in Section S3.

For each layer of the neural network, we extracted the pattern of activations for each image and then computed the pairwise activation pattern dissimilarity for all pairs of images. We then computed the correlation between pattern dissimilarities and actual feature differences (i.e, difference between the actual lengths or brightnesses of the rectangular bars in the two images). We computed this correlation for both absolute (denoted by r_abs_) as well as for relative feature differences (denoted as r_rel_). A positive value for the difference between the two correlation coefficients (r_rel_ – r_abs_) indicated that the Weber’s law was present in a given layer of the neural network.

We also analysed deep networks for the presence of Weber’s law for image intensity, but found highly inconsistent and variable effects. Specifically, the pre-trained VGG-16 network showed Weber’s law for low image intensity levels but not for high intensity levels.

### Experiment 7: Relative Size

We used the stimuli used in a previous study to test whether units in the VGG-16 network encode relative size (Vighneshvel and Arun, 2015). This stimulus set consisted of 24 tetrads. A sample tetrad is shown in Figure 2D, with the stimuli arranged such that images that elicit similar activity are closer. The full stimulus set is shown in Section S3.

In our previous study (Vighneshvel and Arun, 2015), only a small fraction of neurons (around 7%) encoded relative size. To identify similar neurons in the deep network, we first identified the visually responsive units by taking all units with a non-zero variance across the stimuli. For each unit and each tetrad, we calculated a measure of size interactions of the form abs(r11 + r22 – r12 – r21), where r11 is the response to both parts at size 1, r12 is the response to part 1 at size 1 and part 2 at size 2 etc. We then selected the top 7% of all tetrads with the largest interaction effect. Note that this step of selection does not guarantee the direction of the relative size effect. For the selected tetrads, we calculated the relative size index, defined as 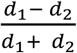 where *d*_1_ and *d*_2_ are distances between the incongruent and congruent stimuli respectively.

### Experiment 8: Decouple pattern shape from surface shape

The stimuli consisted of six patterns superimposed on four surfaces. Each pattern-surface pair was used to create a tetrad of stimuli as depicted in Figure 4E. The full stimulus set consisted of 24 tetrads, which were a subset of those tested in our previous study (Ratan Murty and Arun, 2017). The full stimulus set is shown in Section S3.

In each VGG-16 layer, we selected visually responsive neurons and normalized their responses across all stimuli as before in Experiment 5. We then selected the top 9% of all tetrads with an interaction effect calculated as before, as with the previous study (Ratan Murty and Arun, 2017). For all the selected tetrads we calculated the surface invariance index, defined as 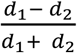 where *d*_1_ and *d*_2_ are distances between incongruent and congruent stimuli.

### Experiment 9: 3D processing

We investigated 3D processing in the VGG-16 network by comparing line drawing stimuli used in a previous perceptual study (Enns and Rensink, 1991). We compared three pairs of shapes: cuboid, cube and frustum of square in isometric view with the corresponding Y junctions. The full stimulus set is shown in Section S3. For each shape, we calculated three distances between equivalent shape pairs with the same feature difference (Figure 5A). We calculated a 3D processing index, defined as 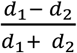 Where *d*_1_ and *d*_2_ are distances between the 3D shape and control conditions respectively.

### Experiment 10: Occlusions

We recreated the stimulus set used in a previous study (Rensink and Enns, 1998) as depicted in Figure 5C. The full stimulus set is shown in Section S3. As before we compared the distance between two pairs of shapes: a pair that differed in occlusion status (occluded vs unoccluded, or two images that differed in their order of occlusion), and an equivalent 2D feature control containing the same feature difference. We then calculated an occlusion index defined as 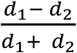 where *d*_1_ and *d*_2_ are the distances between the occluded and control conditions respectively.

### Experiment 11: Object Parts

We performed two experiments to investigate part processing in deep networks. In Experiment 11A, we tested what happens when a break is introduced into an object at a natural cut or an unnatural cut (Xu and Singh, 2002). The full stimulus set is shown in Section S3. The critical comparison is shown in Figure 6A. For each layer of the CNN, we extracted features for the three objects and computed the distance of the intact object with each of the broken objects (d_n_ and d_u_ denote distances to the broken objects with natural and unnatural parts respectively). We then computed a part processing index, defined as 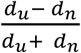.

In Experiment 11B, we asked whether whole object dissimilarities computed on CNN feature representations could be understood as a linear combination of dissimilarities between their natural or unnatural part decompositions as reported previously for visual search (Pramod and Arun, 2016b). We considered seven whole objects that could be broken down into either natural or unnatural parts and recombined the parts to form other objects. That is, we created two sets each containing 49 objects made either from natural or unnatural parts of the original seven objects. The full stimulus set is shown in Section S3. We then selected 492 pairs of objects from each set (including all 21 pairs from the common set) and calculated the feature distances from each layer of the CNN. We fit a part summation model to explain pairwise whole-object distances as a function of pairwise part relations, as described previously (Pramod and Arun, 2016b). We then compared model performance on the 21 pairwise distances between the common objects. We denoted by r_natural_ the correlation between observed and predicted distances assuming natural part decomposition and by r_unnatural_ the model correlation assuming unnatural part decomposition. The natural part advantage was computed as (r_natural_ – r_unnatural_). The same measure was computed for human perception.

### Experiment 12: Global Shape Advantage

We created a set of 49 hierarchical stimuli by combining seven shapes at global scale and the same seven shapes at the local scale (Jacob and Arun, 2019). The full stimulus set is shown in Section S3. We extracted features from all layers of CNNs and calculated the Euclidean distance between all pairs of hierarchical shapes. We calculated the global distance as the mean distance between image pairs having only global change. Similarly, we calculated the local distance as the mean distance between image pairs having only local change. A sample global/local change pair is shown in Figure 6E. We calculated a global advantage index as 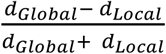.

## ACKNOWLEDGEMENTS

### Funding

This work was supported by the Wellcome Trust-DBT India Alliance Fellowship (Grant # IA/S/17/1/503081) awarded to SPA, and by travel grants to GJ from the Department of Science and Technology, Government of India and from the Pratiksha Trust.

### Author Contributions

GJ, PRT, HK & SPA designed the study, GJ performed Experiments 5,7,8-12 and replicated the analyses for all experiments, HK & GJ performed Experiments 1-3, PRT performed Experiments 4, 6 & 11, and GJ & SPA wrote the manuscript with inputs from PRT & HK.

### Conflicts of interest

The authors declare no conflicts of interest.

## SUPPLEMENTARY MATERIAL

### SECTION S1. RESULTS WITH OTHER FEED FORWARD NETWORKS

The results in the main text were based on testing a specific feedforward network, namely VGG-16. Here, we investigated other feedforward network architectures for the presence of the same perceptual and neural phenomena. We did not test the recurrent networks since unfolding recurrent networks over time make them equivalent to a deep feedforward network (LeCun, Bengio and Hinton, 2015; Liang and Hu, 2015).

#### Methods

We selected four popular pre-trained feedforward networks, all trained on the ImageNet ILSVRC challenge data (Deng *et al*., 2009; Russakovsky *et al*., 2014). We selected architectures that are shallower and deeper than VGG-16, to investigate whether the depth of the network influences the emergence of the perceptual and neural properties. All networks were implemented using MatConvNet framework in MATLAB, and their performance is summarized in Table S1.

*Network 1: AlexNet.* This network won the ILSVRC 2012 challenge by a large margin (Krizhevsky, Alex, Ilya Sutskever, 2012). The network consists of five convolutional layers and three fully-connected layers. Drop-out technique is used fully connected layers to reduce overfitting. The architecture of this network is shallower compared to VGG-16.

*Network 2: GoogLeNet.* This network follows the inception architecture which is well known for better utilization of computing resources inside the network. This network won the classification track of the ILSVRC 2014 challenge (Szegedy *et al*., 2015).

*Network 3: ResNet-50.* ResNet-50 is a shallower variant of the ResNet-152 detailed below.

*Network 4: ResNet-152.* The network uses a residual learning principle which make them capable of training deeper networks without the problem of vanishing gradients (He *et al*., 2016). The ResNet architecture won three tracks (classification, detection and localization) of the ILSVRC 2015 challenge and two tracks (detection and segmentation) of the COCO 2015 challenge.

**Table S1.** Performance of deep networks on the ILSVRC 2012 validation dataset (accuracy reported from MatConvNet website, accessed on 27^th^ November 2019).

| Name of network | Performance |  |
| --- | --- | --- |
|  | Top-1 error (%) | Top-5 error (%) |
| AlexNet | 42.6 | 19.6 |
| VGG-16 | 28.5 | 9.9 |
| GoogLeNet | 34.2 | 12.9 |
| ResNet-50 | 24.6 | 7.7 |
| ResNet-152 | 23.0 | 6.7 |

#### Results

*Experiment 1: Thatcher effect.* The Thatcher index for each network across layers is shown in Figure S1A. It can be seen that the Thatcher index is negative for all networks in their final layers except for GoogLeNet which showed a small positive level in the final layers. For the networks with higher classification performance (GoogLeNet, ResNet-50, ResNet-152), we observed an interesting pattern whereby the Thatcher index is positive in the intermediate layers. This is true even for the VGG-16 network (Figure 2B).

*Experiment 2: Mirror Confusion.* The mirror confusion index for each network is shown in Figure S1B. All networks exhibited an increasing mirror confusion index across layers, just as we observed for VGG-16 (Figure 2D).

*Experiment 3: Scene incongruence.* The classification accuracy for objects in congruent and incongruent scenes is shown for each network in Figure S1C. It can be seen that the deeper architectures show smaller incongruence effects.

*Experiment 4: Multiple object normalization.* The normalization slope for pairs and triplets for all networks is shown in Figure S1C. It can be seen that there is increased normalization in the later layers in all networks.

*Experiment 5: Selectivity across multiple dimensions.* The correlation between sparseness of units in each layer for textures and shapes is shown in Figure S1E. It can be seen that all networks show an increasing trend in later layers. We obtained qualitatively similar results for comparing sparseness on the reference shape set and morph lines (not shown for brevity).

*Experiment 6. Weber’s law.* The Weber’s law measure (difference in correlation for relative vs absolute length) for all networks is shown in Figure S1F. It can be seen that the Weber’s law arises in the later layers for all the networks.

*Experiment 7. Relative size.* The relative size effect for each network across layers is shown in Figure S2A. It can be seen that the relative size effect is extremely weak and variable across networks, and never approaches the levels observed in the brain (relative size index = 0.39).

*Experiment 8: Decoupling patterns from surfaces.* The surface invariance index for each network across layers is shown in Figure S2B. It can be seen that the index is consistently negative for all networks, as observed for VGG-16.

*Experiment 9: 3D processing.* The 3D processing indices (for Condition 1 & 2) for each network across layers is shown in Figure S2C. It can be seen that the 3D processing indices are generally negative for all networks, and even if the index is positive, the levels are much smaller than observed in humans.

*Experiment 10: Occlusions.* The occlusion indices for each network across layers is shown in Figure S2D. It can be seen that both indices are consistently negative across layers, as observed for VGG-16.

*Experiment 11: Object parts.* The natural part advantage for Experiment 11B is shown for each network across layers in Figure S2E. It can be seen that the natural part advantage is highly variable across networks, with GoogLeNet showing levels comparable to humans in the later layers.

*Experiment 12: Global shape advantage.* The global advantage index for each network across layers is shown in Figure S2F. Across all networks, there is a slight global advantage in the intermediate layers, which reverses into a local advantage in the later layers. Thus, it appears that all the feedforward networks are using local features for classification.

**Figure S1:**
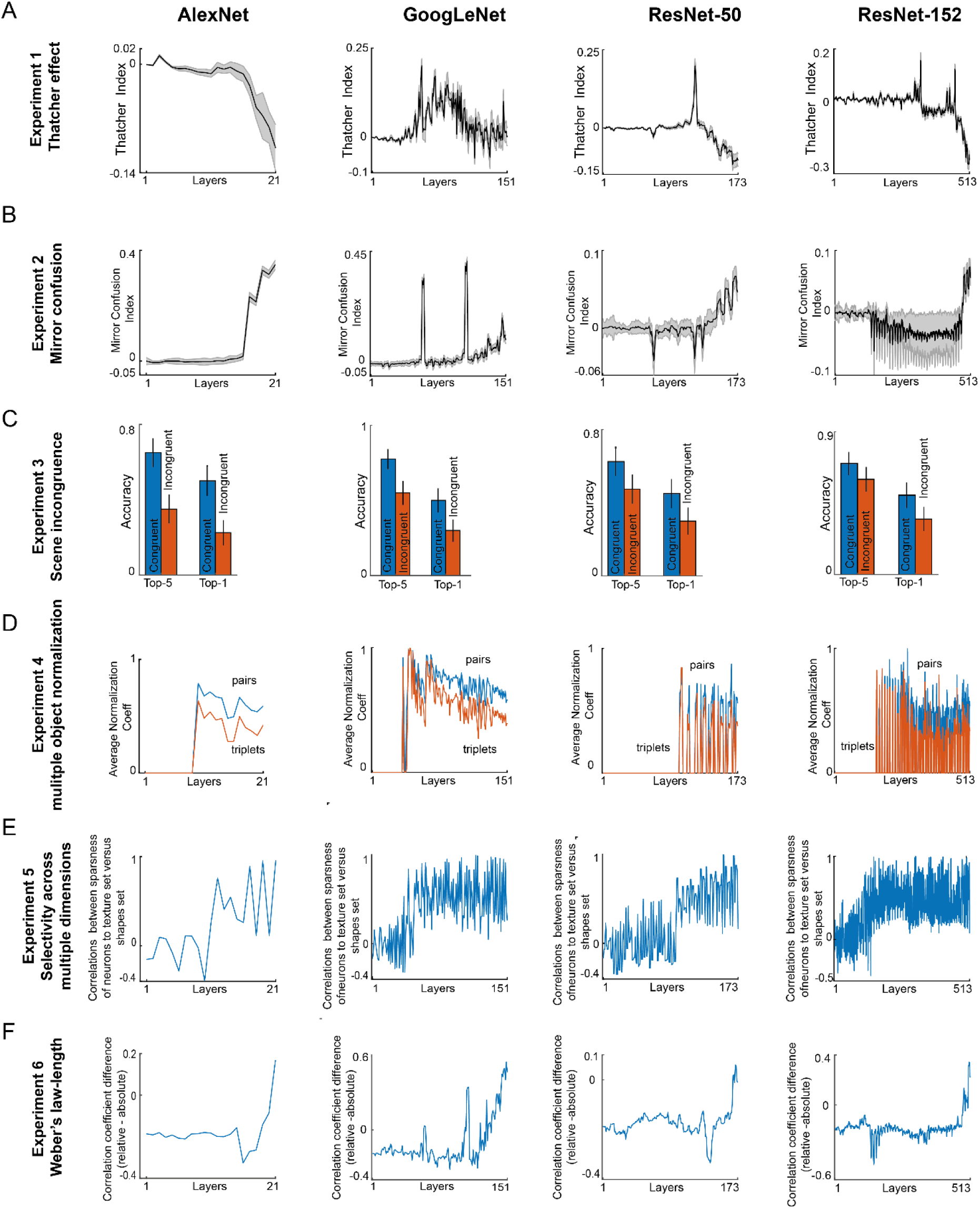
Experiments 1-6 with other feed-forward networks. Each column represents a deep network (from left to right: AlexNet, GoogLeNet, ResNet-50 and Reset-152) and each row represents an experiment (A-F: experiment 1-6). In each subplot an experiment specific index or measure is plotted across layers.

**Figure S2:**
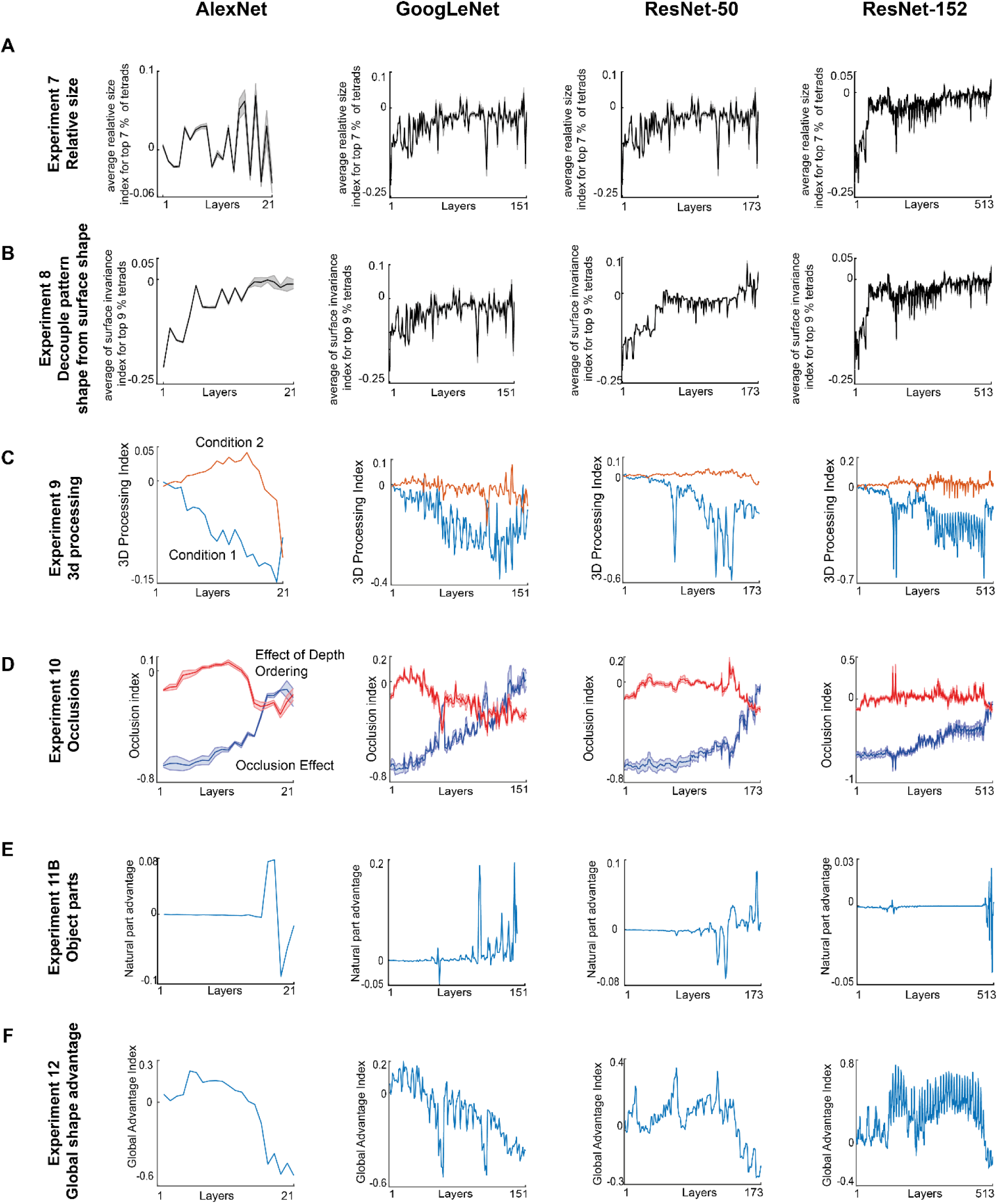
Experiments 7-12 for other feedforward networks. All conventions as in Figure S1.

### SECTION S2. RESULTS WITH RANDOMLY INITIALIZED NETWORKS

We have shown that deep neural networks trained for object-recognition show some perceptual phenomena but not others, but we wondered whether any of these phenomena can be observed even in an untrained network, simply as a consequence of the architecture. To address this issue, we repeated Experiments 1-12 on an untrained VGG-16 architecture with randomly initialized weights.

#### Methods

We estimated the probability distribution functions of the weights in each layer of the pre-trained VGG-16 network and picked the weights randomly from this estimated distribution. This process was done layer-wise to obtain the randomly initialized network. We then performed all Experiments on this network except for Experiment 3 (scene incongruence) since this experiment requires object classification. We then performed all data analyses exactly as before.

#### Results

The results for the random network and the pre-trained VGG-16 network are shown together in Figure S3. In most cases (Experiments 1, 2, 6-11), we observed no specific trend for the random network towards or away from human/neural levels, which is what would be expected since the random network has no specific bias or training. We observed systematic and interesting differences for the other experiments, which we discuss in detail below.

*Experiment 4: Multiple Object normalization.* The divisive normalization slope for the random network is shown in Figure S3C-D. Here, the random network showed perfect divisive normalization, in that the net response to AB is exactly the average of the responses to A & B separately. To investigate this puzzling observation further, we looked at the unit activations in the random network. We found that the activations for any pair of natural images was highly correlated (correlation of layer-37, mean ± sd: r = 0.98 ± 0.01), suggesting that these units were not very selective for images. This means that every image activates the network in the same way. As a result, the response to AB and the response to A & B separately would be identical, giving rise to a perfect slope of 0.5 in the relationship between the response AB and the sum of responses A + B. Thus, the divisive normalization observed in the random network is a trivial outcome of its lack of image selectivity.

*Experiment 5: Selectivity across multiple dimensions.* Here too the random network shows a high correlation between shape and texture selectivity (Figure S3E). We suspect that this too is a consequence of the very low selectivity for images in the random network, whereby some units have zero selectivity (and therefore respond equally to all images) and others have weak selectivity (and therefore show slight differences in the response across images).

*Experiment 12: Global shape advantage*. Here we observed an interesting pattern: the random network showed a global advantage that increased across layers. This is likely due to increased pooling in the higher layers, but interestingly the pre-trained VGG-16 network shows the opposite pattern. Thus, it appears that object classification training abolishes the global advantage that is intrinsic to the network architecture. We speculate that this local advantage might arise because of the demands of distinguishing between highly similar categories present in the ImageNet dataset (e.g. there are 90 categories of dogs among the total of 1000 categories in ImageNet). Testing this possibility will require training the VGG-16 architecture on highly distinctive object classes.

**Figure S3.**
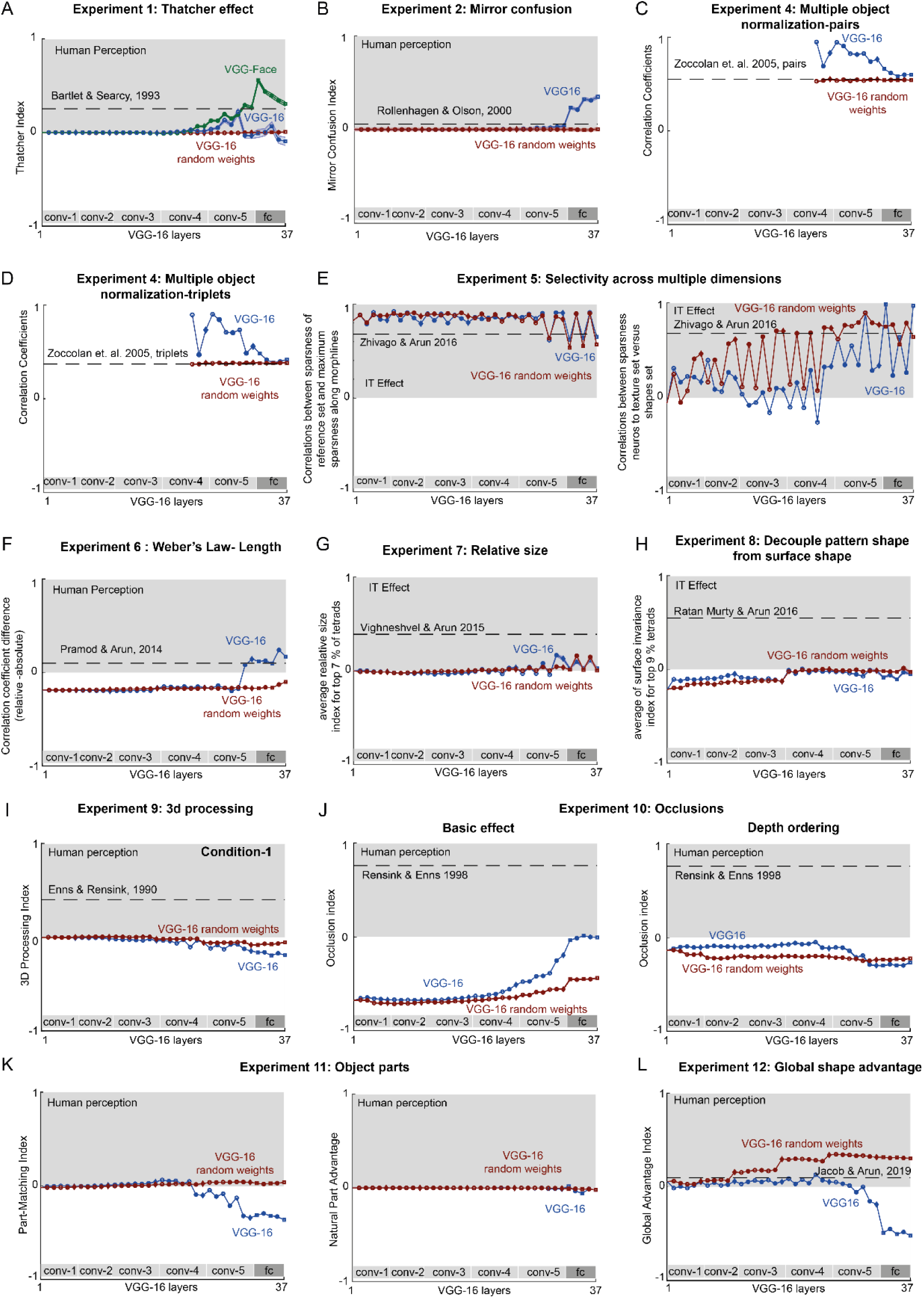
Experiments 1-12 for randomly initialized networks. Results for Experiments 1-12 (except for #3) are shown. In each panel, the corresponding experiment-specific index is plotted for the randomly initialized VGG-16 (*red*) with the pre-trained VGG-16 (*blue*). All other conventions are as in the main text.

### SECTION S3. STIMULI USED IN EXPERIMENTS 1-12

**Figure S4.**
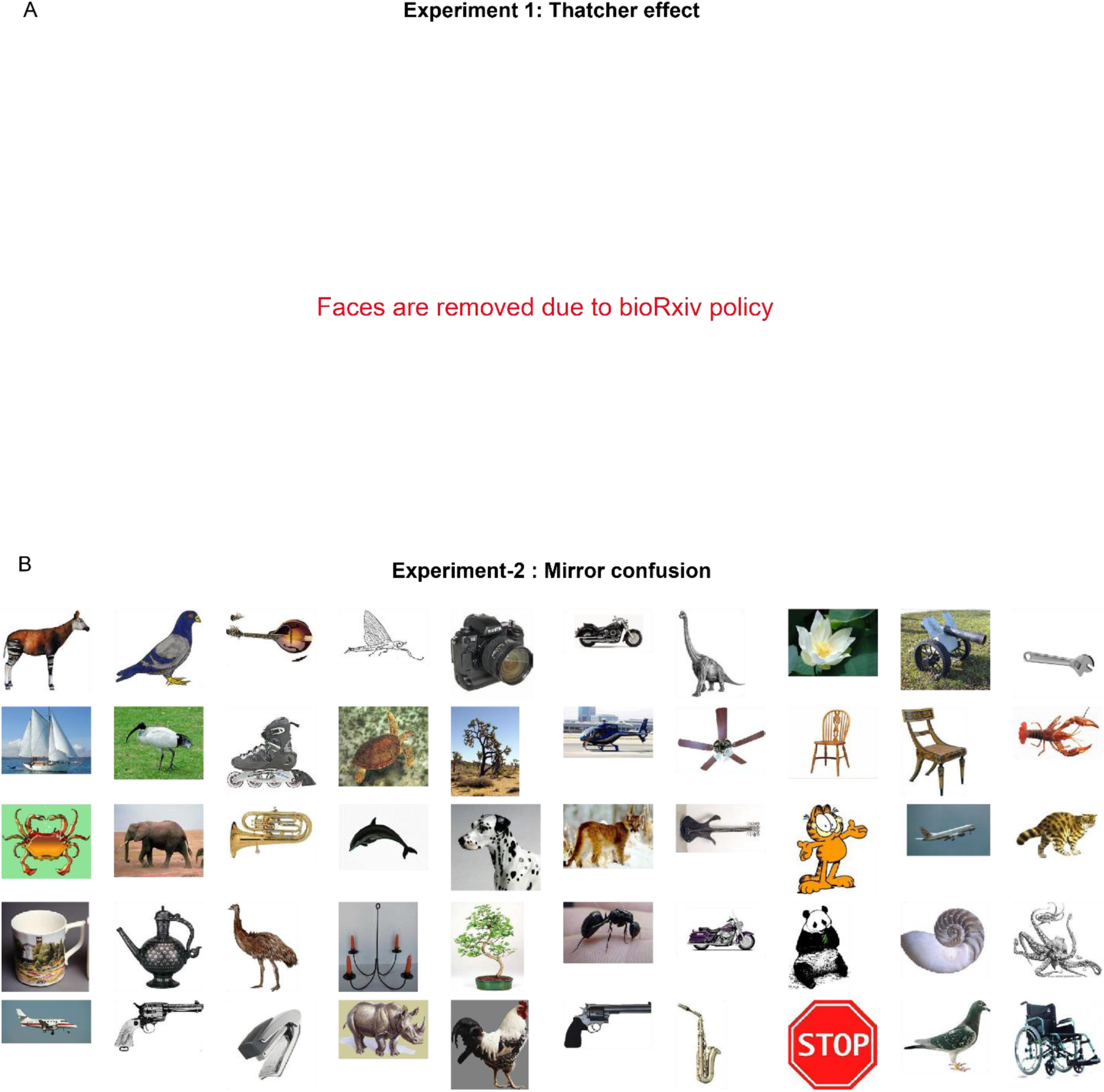
Stimulus set used for Experiments 1 & 2.

**Figure S5.**
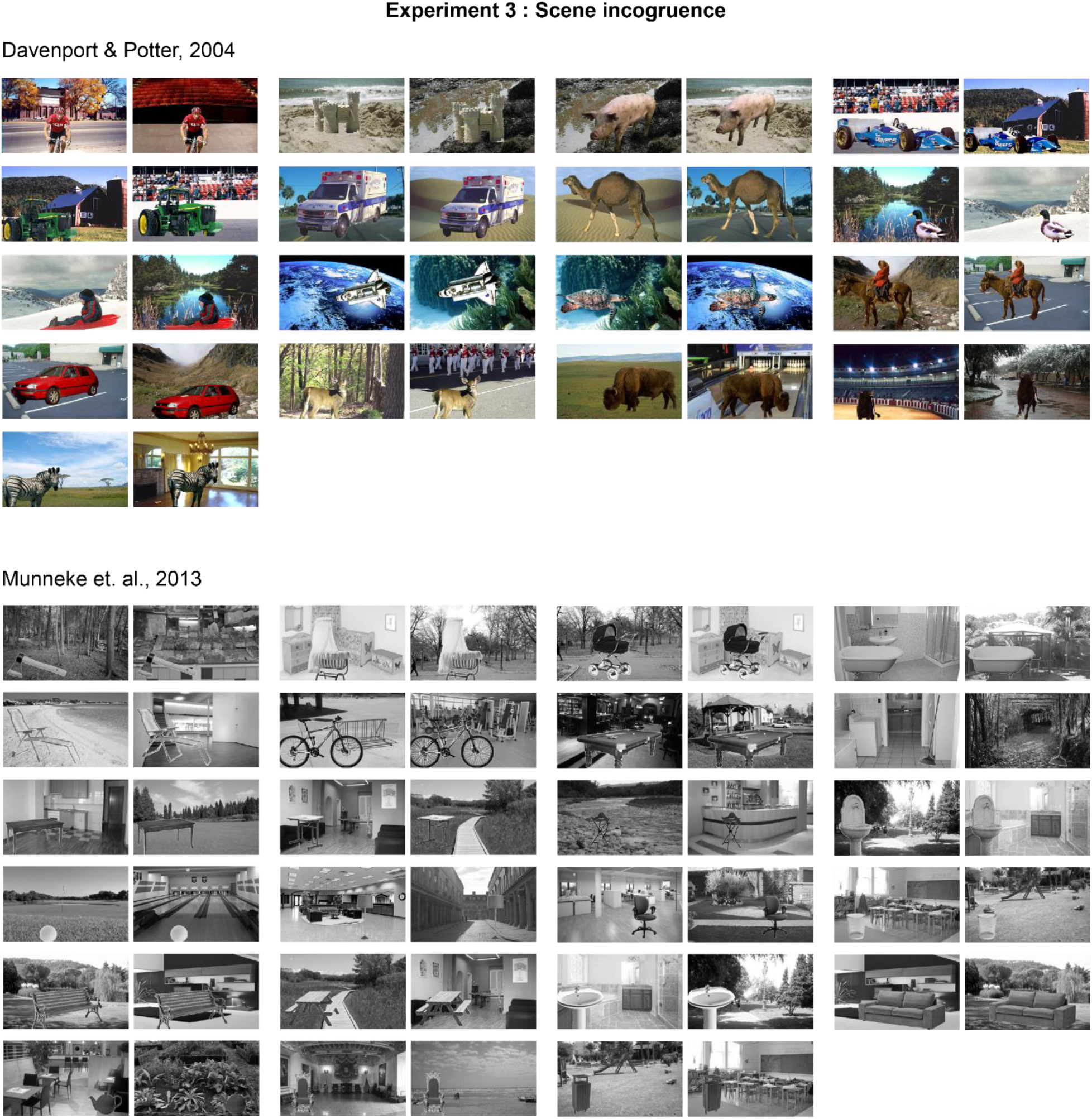
Stimulus set used for Experiment 3. Each pair of images depicts an object against a congruent and incongruent background. Stimulus set reproduced with consent.

**Figure S6.**
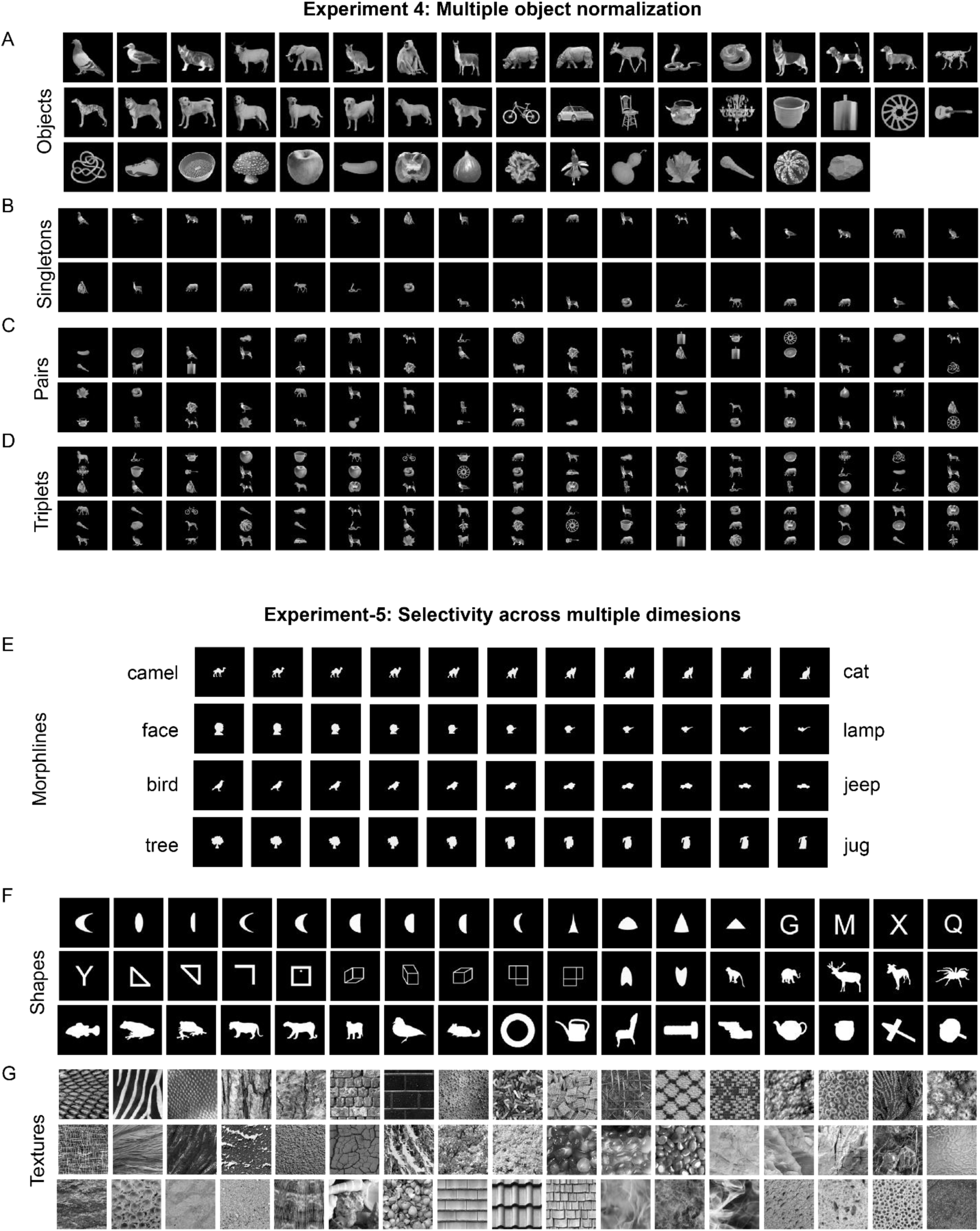
Stimulus set used for Experiments 4 & 5. Plots A-D shows the images used for experiment 4 and plots E-G shows the images used in experiment-5. A) 49 natural images used for experiment-4 B) Singletons are formed by placing the 49 objects either at top, middle or bottom. 34 selected singletons are shown from 147 singletons used in this experiment. C) 34 selected pairs are shown from 200 pairs used in experiment-4. D) 34 selected triplets are shown from 200 triplets used in experiment-4. E) Shows all four morphlines used in experiment-5. F) Shown 51 selected shapes of the total set of 120 shapes. G) Shown 51 selected textures out of the total set of 120 textures.

**Figure S7.**
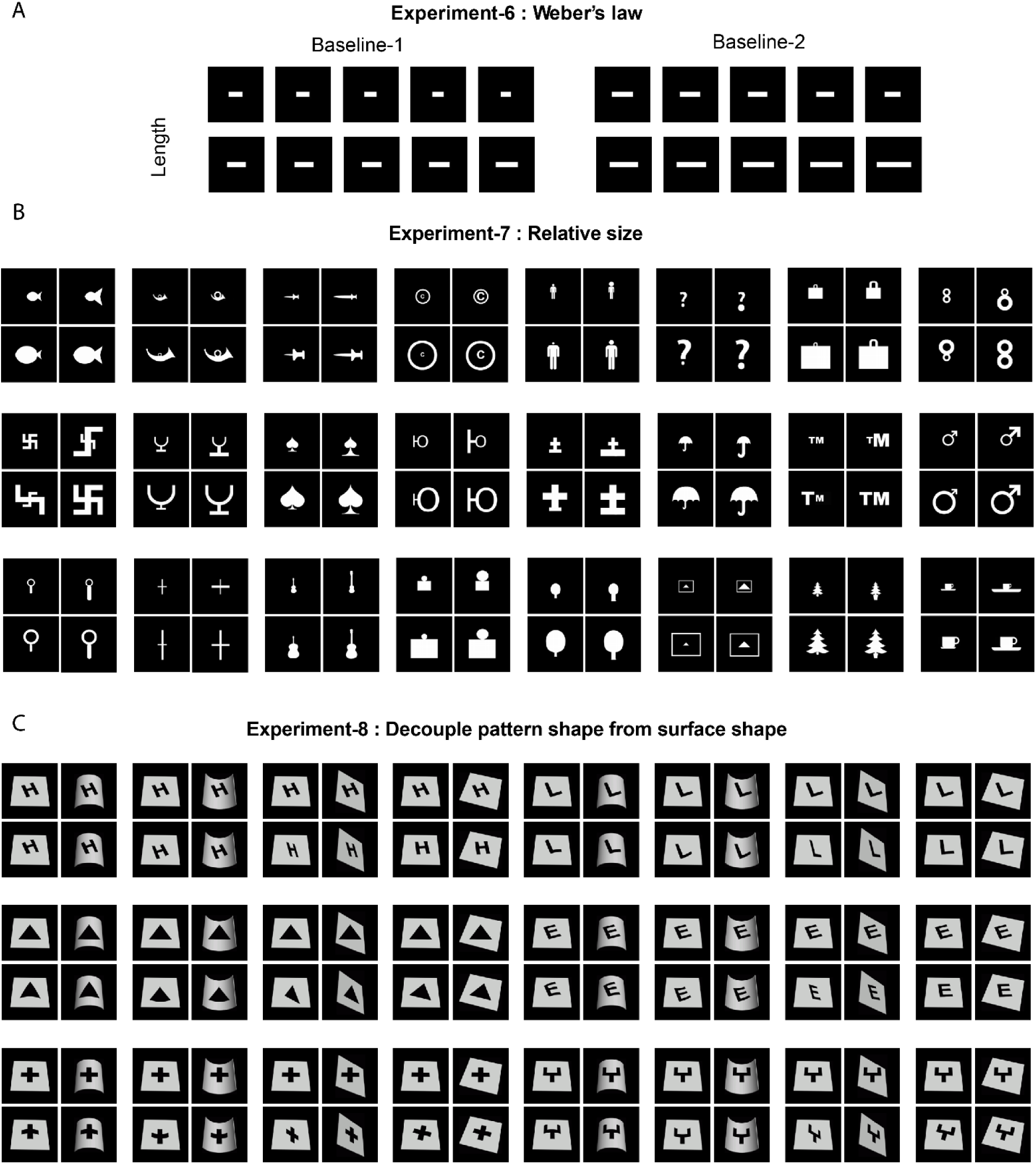
Stimulus set used for Experiments 6-8. A) Shows the images used in Weber’s law experiment. There are two baselines and each column is an image pairs which has equal length difference from the baseline. B) Shows 24 tetrads used in Relative Size experiment. C) Shows 24 tetrad used in surface invariance experiments. Tetrads are made by transforming eight shapes onto five different surfaces.

**Figure S8.**
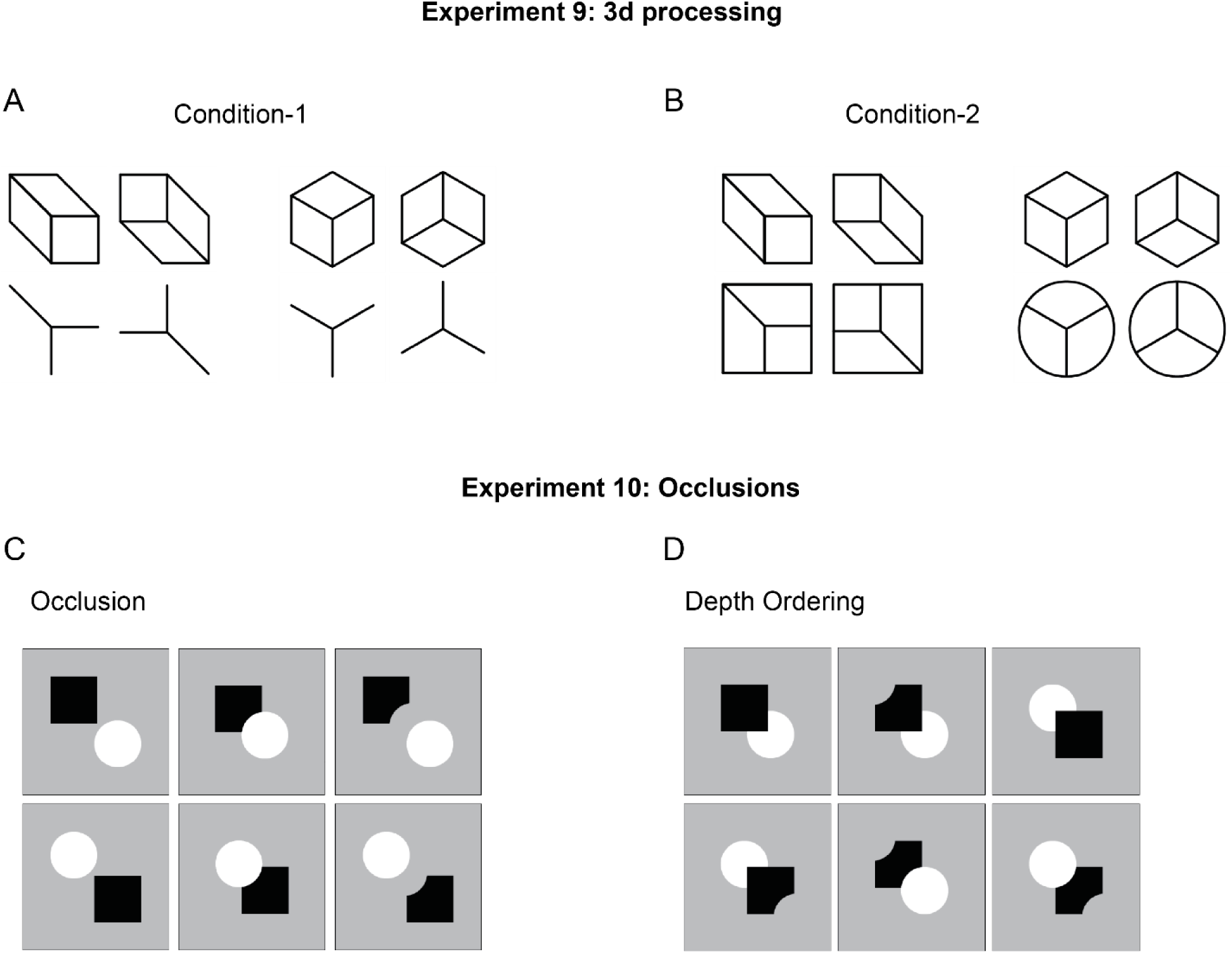
Stimulus set used for Experiments 9-10. A) Two sets of images used to compare the 3D perception in the CNNs. Images in top row have 3D effect whereas the images in the bottom row have an equivalent same feature difference without a perceived 3D difference. B) Two sets of images used to compare the 3D perception in the CNNs. Images in top row have 3D effect whereas the images in the bottom row has the same feature difference and an additional common outer shape but not the 3D perception. C) Each row shows a set of images used for testing basic occlusion effect. D) Each row shows a set of images used for testing depth ordering effect.

**Figure S9.**
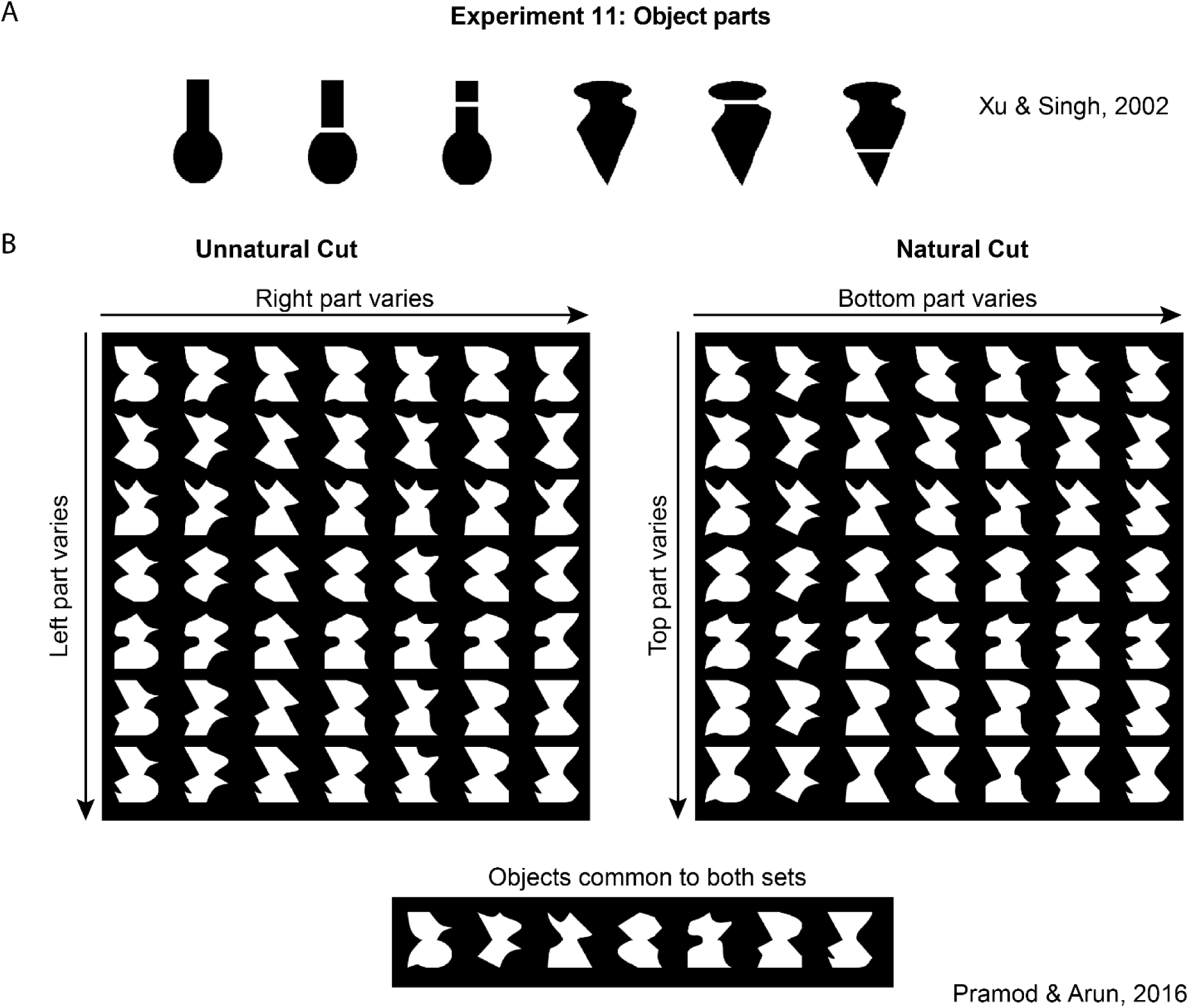
Stimulus set used for Experiments 11. A) Shows the images used to check part processing in experiment-11 B) Shows the images used to check part advantage in experiment-11. The seven shapes in the diagonal position of both Unnatural and natural set are the same.

**Figure S10.**
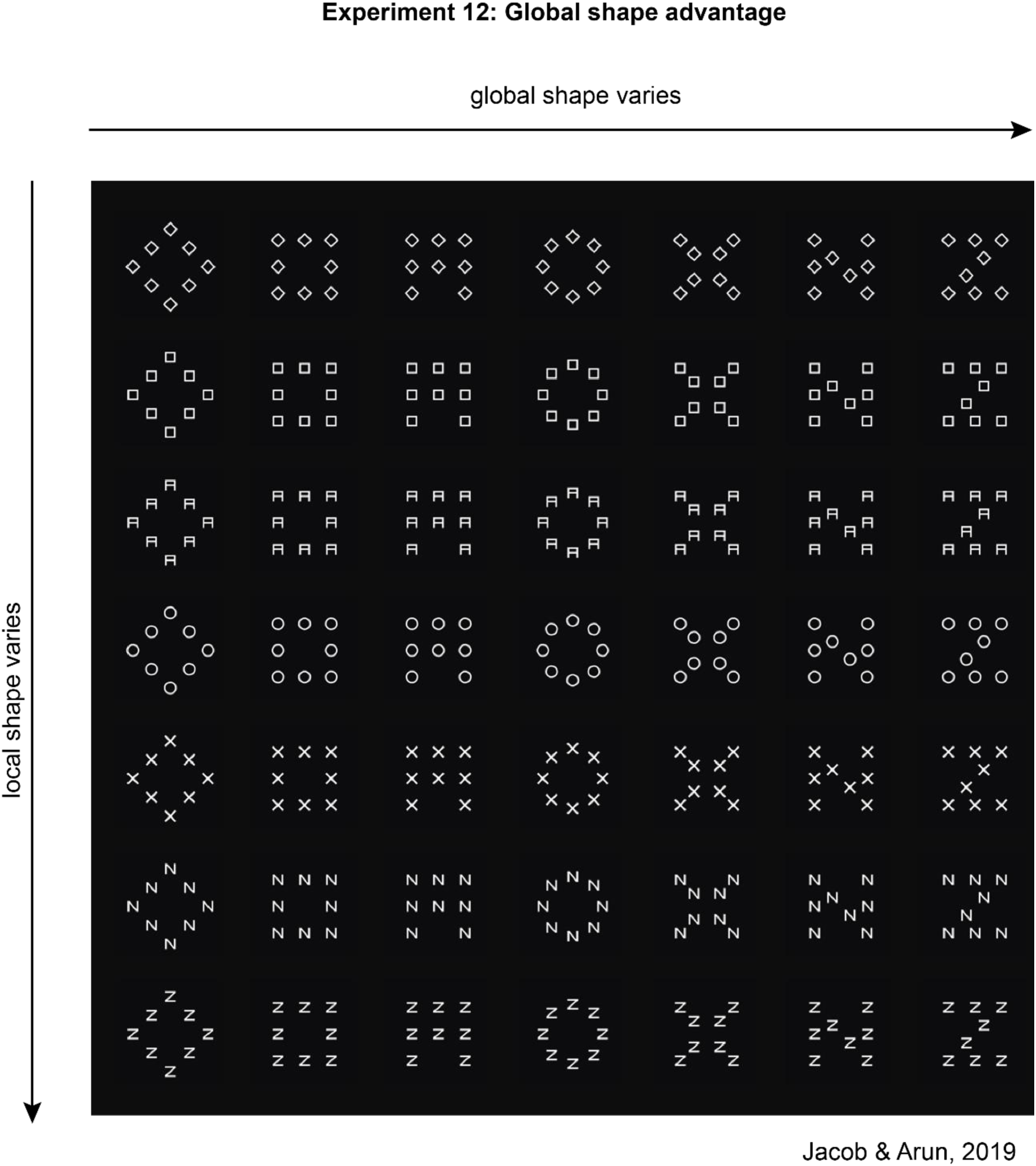
Stimulus set used for Experiments 12. A total of 49 images used to check global advantage. These images are formed by all combinations of seven shapes are global and local scales.

